# Niche-specific microbial community structure of subgingival plaque in periodontitis

**DOI:** 10.64898/2026.08.12.744510

**Authors:** Qingxiu Li, Guangmei Li, Zhiwen Liu, Zhenjun Li

## Abstract

Subgingival biofilms in periodontitis exhibit spatial heterogeneity, yet the organization of microbial communities across periodontal niches remains incompletely defined. Using paired sampling and 16S rRNA gene sequencing, we characterized non-attached and attached subgingival plaque from patients with periodontitis, together with non-attached plaque from periodontally healthy individuals. Across diversity metrics and ordination analyses, non-attached plaque from periodontitis patients occupied positions between healthy-associated and attached-plaque communities. Taxonomically, these communities contained both health-associated commensals and anaerobic genera commonly enriched in periodontitis. Network analysis identified differences in association-network topology among niches, with the non-attached periodontitis network containing more retained associations than the healthy network. These cross-sectional results describe niche-associated patterns of subgingival community composition and association structure. They do not establish temporal progression, direct microbial interactions, or clinical utility.

## Introduction

Periodontitis is a prevalent inflammatory disease and a major cause of tooth loss in adults worldwide [1–3]. It is associated with changes in the subgingival microbiome, including shifts from communities dominated by health-associated taxa toward communities enriched in anaerobic and inflammation-associated taxa [4–6]. Periodontitis is increasingly understood as a community-level condition involving coordinated changes in microbial composition within subgingival biofilms.

The subgingival environment is heterogeneous, with gradients in oxygen availability, nutrient sources, and host-derived inflammatory factors [8–10]. This spatial structure supports distinct microbial communities within the periodontal pocket [11,12]. Previous sequencing studies have often contrasted periodontal health and disease while treating subgingival plaque as a single ecological entity [13,14]. Different plaque compartments within the same periodontal site may nevertheless harbor distinct microbial assemblages.

Within periodontal pockets, non-attached plaque resides in the pocket lumen, whereas attached plaque is associated with the root surface [15,16]. These plaque types differ in physical structure, exposure to shear forces, and proximity to host tissues [17]. The microbial organization of non-attached plaque remains less well characterized than that of attached plaque. It is therefore unclear whether non-attached plaque is simply a diluted form of attached-plaque communities or a distinct configuration with its own compositional and association-network features.

We used paired sampling and 16S rRNA gene sequencing to characterize the taxonomic composition and community organization of non-attached and attached subgingival plaque from participants with periodontitis, together with non-attached plaque from periodontally healthy individuals. We integrated diversity metrics, compositional analyses, and microbial association networks to describe niche-specific patterns. The study was cross-sectional; all comparisons are therefore interpreted as associations across sampled niches rather than evidence of disease progression, direct interactions, or clinical utility.

## Materials and Methods

### Study population and sample collection

Participants were recruited from the Department of Stomatology, the Second Xiangya Hospital of Central South University. Participants with periodontitis and periodontally healthy controls were diagnosed by two calibrated periodontists using the clinical criteria recorded at enrolment and with reference to the 2017 World Workshop classification [23]. Because stage and grade data were not available in the analysis dataset, diagnoses are reported here as periodontitis rather than as stage- or grade-specific categories. Inclusion criteria included the presence of at least 20 natural teeth and no history of systemic disease or antibiotic use within one month before sampling. Exclusion criteria included pregnancy, orthodontic treatment, betel nut chewing, or refusal to provide informed consent. A total of 46 individuals were enrolled, including 36 patients with periodontitis and 10 periodontally healthy controls. In patients with periodontitis, paired non-attached (CP-NA) and attached (CP-AD) subgingival plaque samples were collected from the deepest periodontal pocket, yielding 36 paired disease-site sets (72 samples). In healthy controls, non-attached plaque (PH-NA) was collected from the mesiobuccal sites of first molars (10 samples). The final dataset therefore comprised 82 samples. Plaque samples were obtained using sterile Gracey curettes, immediately placed on ice, and stored at −80 °C until processing.

All participants provided written informed consent. The study was conducted in accordance with the Declaration of Helsinki and was approved by the Ethics Committee of the Second Xiangya Hospital of Central South University (approval number: 2021–038). Clinical and sampling characteristics are summarized in Table 1.

**Table 1.**
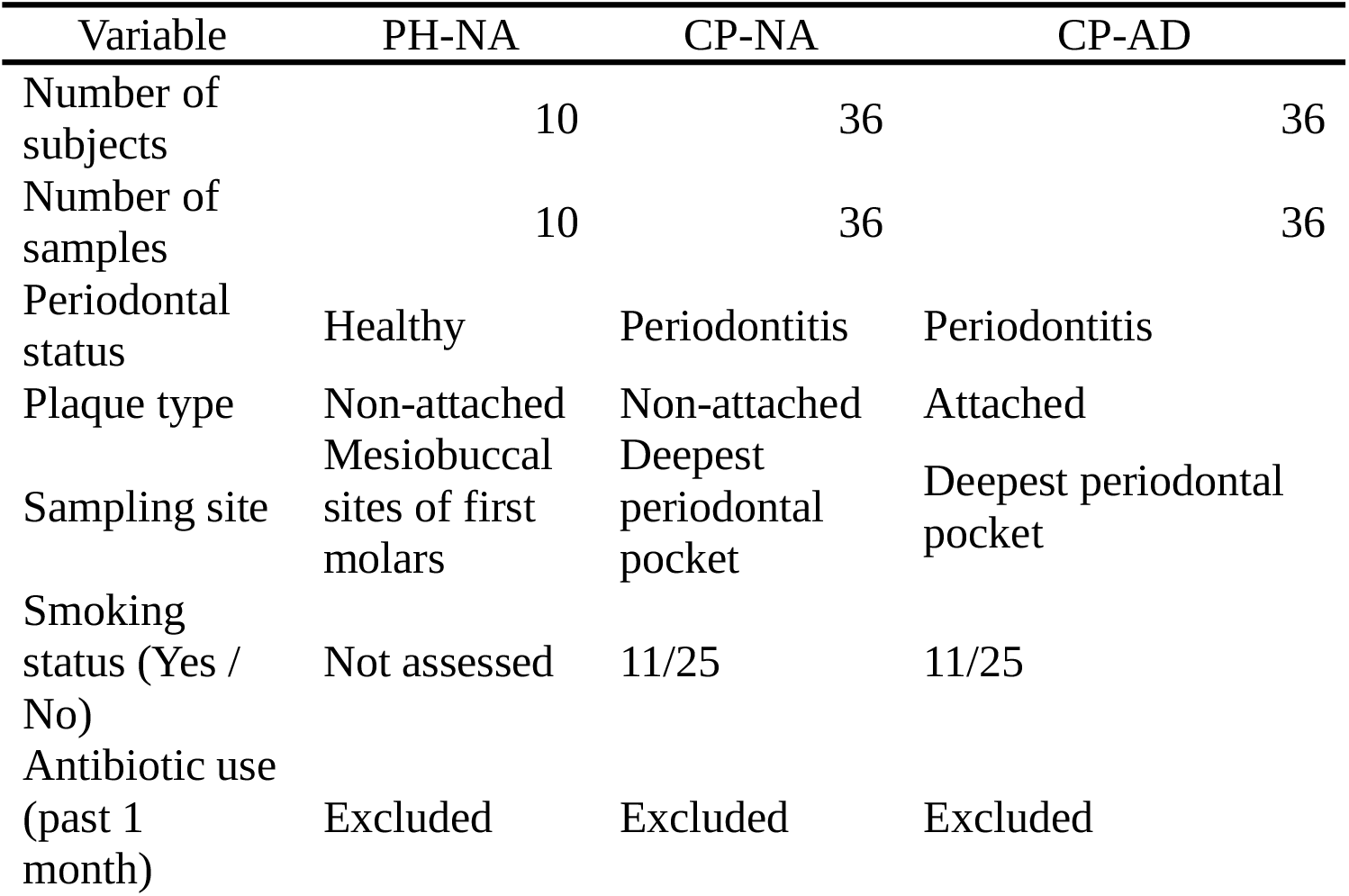
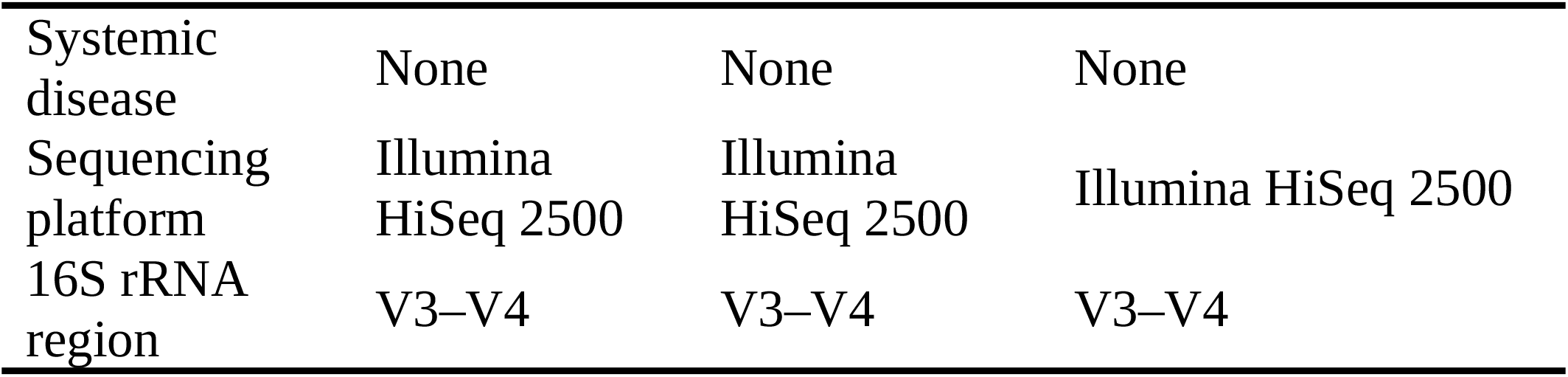
Clinical and sampling characteristics of study participants.

### DNA extraction and 16S rRNA gene sequencing

Genomic DNA was extracted from plaque samples using SDS and proteinase K lysis followed by phenol–chloroform purification. DNA concentration and quality were assessed prior to amplification. The V3–V4 region of the bacterial 16S rRNA gene was amplified using primers 341F and 806R and a high-fidelity DNA polymerase. Amplicons were purified, pooled at equimolar concentrations, and sequenced on an Illumina HiSeq 2500 platform with 2 × 300 bp paired-end reads. Extraction blanks and PCR-negative controls were included to monitor potential contamination [24].

### Sequence processing and taxonomic assignment

Raw sequencing reads were processed using DADA2 (v1.28.0) in R (v4.3.2). Reads were quality filtered, denoised, merged, and screened for chimeras to generate amplicon sequence variants (ASVs) [25]. Taxonomic assignment was performed using the SILVA reference database (release 138.1) with a naïve Bayesian classifier [26,27]. ASVs with low prevalence or low relative abundance were removed to reduce sparsity. For downstream analyses, ASV counts were normalized to relative abundances and aggregated at the genus level.

### Diversity and community composition analyses

Alpha diversity was assessed using Shannon and Chao1 indices calculated from the ASV count matrix [28]. The prespecified PH-NA versus CP-NA comparison was evaluated with a two-sided Wilcoxon rank-sum test; the CP-NA versus CP-AD comparison used a two-sided paired Wilcoxon signed-rank test. Reported alpha-diversity P values are unadjusted, two-sided values for these prespecified comparisons. Beta diversity was calculated using Bray–Curtis dissimilarities and visualized by principal coordinates analysis (PCoA) [29,30]. Community-level differences were assessed using permutational multivariate analysis of variance (PERMANOVA) with 999 permutations, and dispersion was evaluated using betadisper [31].

### Differential abundance analysis

Genus-level relative abundances were compared using Wilcoxon rank-sum tests for independent comparisons (PH-NA versus CP-NA and PH-NA versus CP-AD) and Wilcoxon signed-rank tests for the paired CP-NA versus CP-AD comparison. Effect sizes were estimated as Hodges–Lehmann differences, with 95% confidence intervals; P values were adjusted across genera within each comparison using the Benjamini–Hochberg procedure. The 25 genera with the smallest adjusted P values are displayed in Fig. 3D–F.

### Microbial association network analysis

Genus-level association networks were constructed separately for each niche using Spearman correlations among genera. Genera present in at least 20% of samples within a niche were retained. Pairwise correlations were Benjamini–Hochberg adjusted within each network; edges with |ρ| ≥ 0.5 and adjusted P < 0.05 were retained. Networks were visualized using igraph and ggraph, and node/edge counts, density, mean degree, modularity, and centrality summaries were calculated from the retained networks. Network analyses were descriptive and did not imply direct ecological interactions or causal relationships.

## Results

### 1. Subgingival microbial configurations across periodontal niches

Sequencing of the V3–V4 region of the 16S rRNA gene generated 1,997,033 high-quality reads across all samples, yielding 7,621 amplicon sequence variants (ASVs). After prevalence and abundance filtering, 1,926 ASVs were retained for downstream analyses, representing 11 bacterial phyla and 84 genera. Rarefaction analysis indicated that sequencing depth was sufficient to capture most observed diversity across sample groups (Fig. 1F). At the genus level, subgingival microbial communities differed across sampling niches (Fig. 1). Periodontally healthy non-attached plaque (PH-NA) was dominated by facultative commensal genera, including Streptococcus, Rothia, and Haemophilus. In contrast, plaque from participants with periodontitis showed increased relative abundances of anaerobic genera such as Fusobacterium, Prevotella, Porphyromonas, and Tannerella, with the highest enrichment observed in attached plaque (CP-AD).

**Fig. 1.**
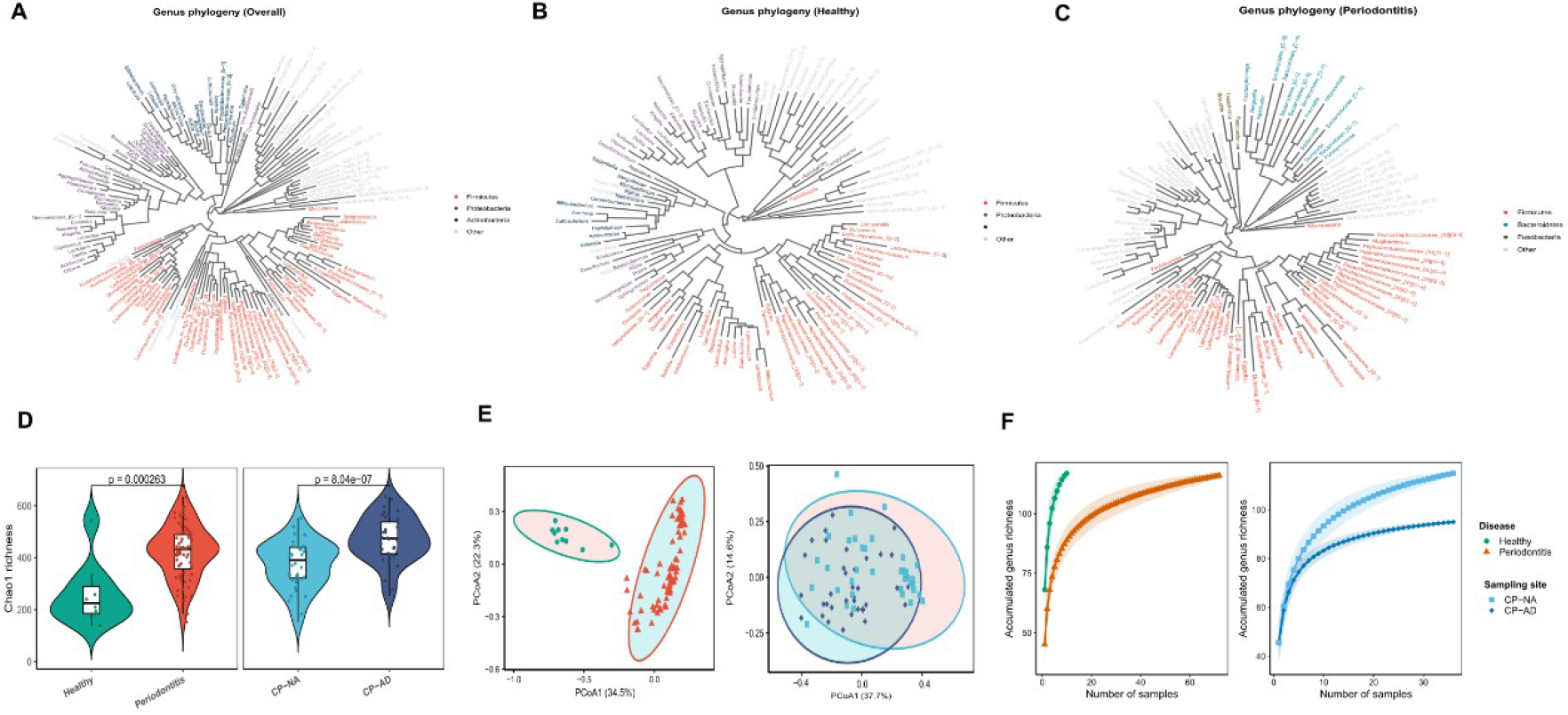
Phylogenetic and diversity characteristics of subgingival microbial communities across periodontal niches. (A) Overall genus-level phylogenetic tree showing the distribution of dominant bacterial phyla across all subgingival samples. (B) Genus-level phylogenetic tree of non-attached plaque from periodontally healthy individuals (PH-NA). (C) Genus-level phylogenetic tree of subgingival plaque from participants with periodontitis, including non-attached (CP-NA) and attached (CP-AD) samples. (D) Violin plots of Shannon and Chao1 diversity indices. (E) Principal coordinate analysis based on Bray–Curtis dissimilarities. (F) Rarefaction curves of observed amplicon sequence variants (ASVs).

Non-attached plaque from participants with periodontitis (CP-NA) displayed a mixed taxonomic composition. CP-NA samples retained health-associated genera while showing increased abundance of anaerobic taxa commonly associated with periodontitis. This pattern distinguished CP-NA from both PH-NA and CP-AD samples. Phylogenetic visualization of dominant genera further illustrated these compositional patterns (Fig. 1A–C).

### 2. Diversity patterns across periodontal niches

Alpha diversity differed among subgingival plaque types (Fig. 1D). Shannon diversity was lower in PH-NA than in CP-NA (Wilcoxon rank-sum P = 2.60 × 10^−6^). Within the paired periodontitis samples, Shannon diversity was lower in CP-NA than in CP-AD (Wilcoxon signed-rank P = 2.57 × 10^−6^).

Chao1 richness showed the same direction of difference (PH-NA versus CP-NA, P = 3.25 × 10^−3^; CP-NA versus CP-AD, paired P = 8.04 × 10^−7^). Summary statistics and design-aware tests are reported in Supplementary Table S1B–F. These cross-sectional differences describe sampling-niche associations and should not be interpreted as evidence of temporal progression.

Beta diversity analysis based on Bray–Curtis dissimilarities showed separation among periodontal niches (Fig. 1E). PCoA revealed distinct clustering of PH-NA and CP-AD samples, with CP-NA samples occupying intermediate positions in ordination space. In a non-paired comparison, PH-NA and CP-NA differed by PERMANOVA (R^2^ = 0.347, P = 0.001). In the paired CP-NA versus CP-AD comparison, restricted within-participant label permutations also supported a difference (R^2^ = 0.081, P = 0.001). Dispersion differed in both comparisons (P = 0.020 and P < 0.001, respectively); therefore, these PERMANOVA findings should be interpreted as overall distributional differences rather than solely as differences in group centroids. Design-aware PERMANOVA and dispersion results are reported in Supplementary Table S1D.

### 3. Taxonomic features of non-attached plaque in periodontitis

Phylum-level profiles revealed systematic compositional differences across periodontal niches (Fig. 2A). PH-NA communities were enriched in Proteobacteria and Actinobacteria, whereas CP-AD samples showed increased relative abundances of Bacteroidetes, Fusobacteria, Spirochaetes, and Synergistetes. CP-NA samples exhibited phylum-level compositions that were intermediate between PH-NA and CP-AD, with elevated representation of anaerobic phyla compared with healthy controls but lower enrichment than attached plaque. At the genus level, hierarchical clustering based on relative abundance highlighted niche-specific patterns (Fig. 2B). Health-associated genera clustered predominantly with PH-NA samples, whereas lesion-associated anaerobes were concentrated in CP-AD samples. CP-NA samples showed mixed clustering patterns, reflecting heterogeneity in genus-level abundance profiles.

**Fig. 2.**
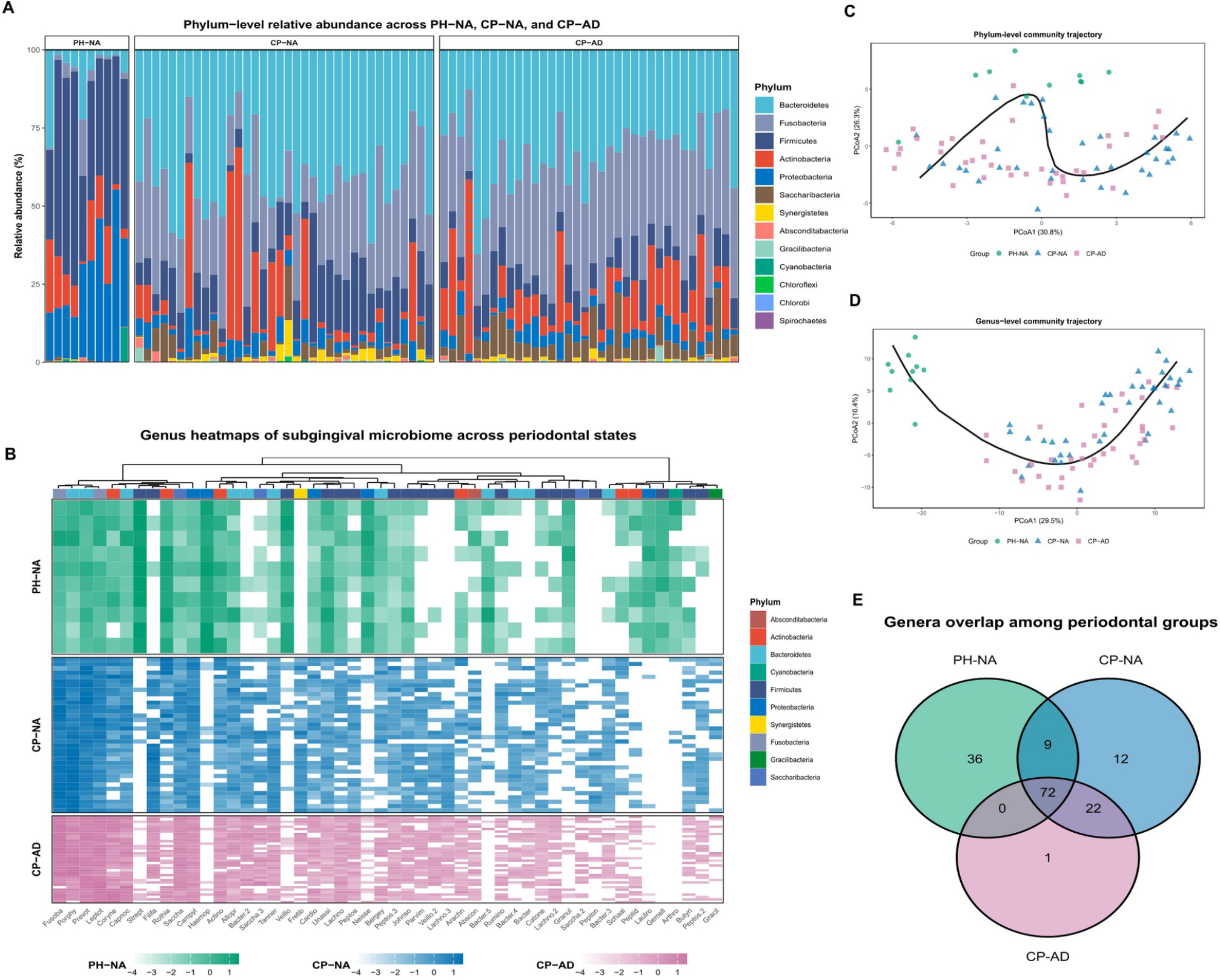
Taxonomic composition and community structure of subgingival microbiota across periodontal niches. Curves are descriptive and do not represent longitudinal trajectories. (A) Stacked bar plots showing phylum-level relative abundance of subgingival microbiota in PH-NA, CP-NA, and CP-AD groups. (B) Hierarchical clustered heatmap of genus-level relative abundances across periodontal niches; color intensity indicates z-score–normalized abundance. (C) Principal coordinate analysis (PCoA) based on Bray–Curtis dissimilarities at the phylum level, showing niche-specific separation of microbial communities. (D) PCoA at the genus level illustrating differences in community composition among periodontal niches. (E) Venn diagram depicting shared and niche-associated genera among PH-NA, CP-NA, and CP-AD groups.

Ordination analyses at both the phylum and genus levels supported these observations (Fig. 2C–D). CP-NA samples consistently occupied positions between PH-NA and CP-AD clusters, indicating compositional differences that were reproducible across multiple analytical resolutions. Analysis of shared genera revealed a conserved core microbiota comprising 72 genera present across all niches, collectively accounting for more than 85% of total relative abundance (Fig. 2E). Beyond this shared core, each niche displayed distinct expansions of accessory taxa, contributing to niche-specific community profiles.

### 4. Niche-specific microbial association networks

Genus-level microbial association networks based on Spearman correlations revealed distinct patterns of community organization across periodontal niches (Fig. 3A–C). The PH-NA network comprised a moderate number of nodes and edges and exhibited a relatively high degree of modularity, with associations distributed across multiple modules and no single genus dominating network connectivity.

**Fig. 3.**
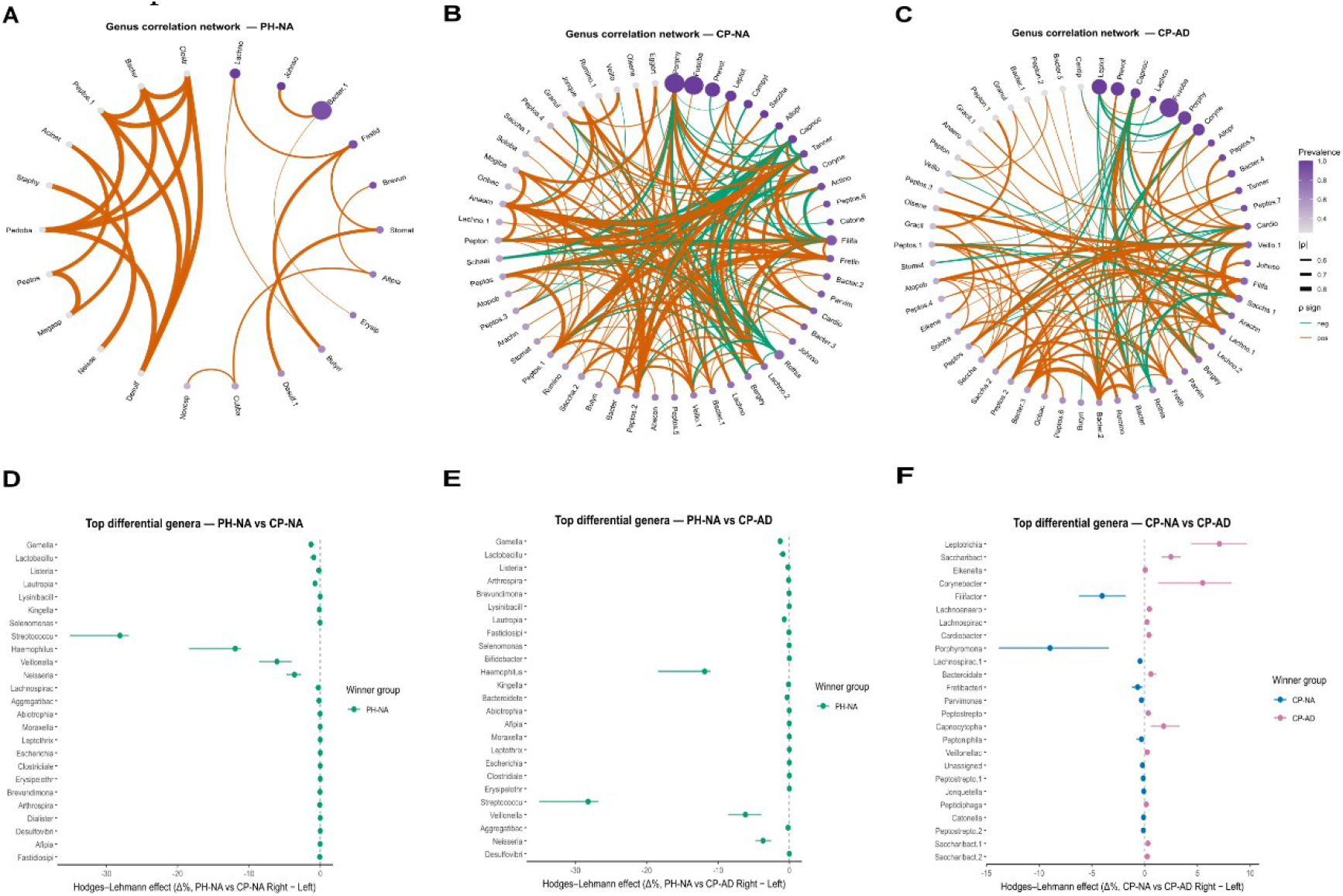
Microbial association networks and differential taxa across periodontal niches. Differential-abundance panels display the 25 genera with the smallest Benjamini–Hochberg-adjusted P values in each comparison. (A–C) Genus-level association networks based on Spearman correlations for (A) periodontally healthy non-attached plaque (PH-NA), (B) non-attached plaque from periodontitis (CP-NA), and (C) attached plaque from periodontitis (CP-AD). Node size represents genus prevalence, and edge width and color indicate correlation strength and sign (orange, positive; green, negative). (D–F) Forest plots showing genera differentially abundant between periodontal niches based on Hodges–Lehmann effect size estimation: (D) PH-NA vs CP-NA, (E) PH-NA vs CP-AD, and (F) CP-NA vs CP-AD. Points indicate median effect sizes, horizontal bars represent 95% confidence intervals, and colors denote the group with higher relative abundance.

Compared with PH-NA, the CP-NA network contained more retained edges and had higher density and mean node degree (Supplementary Table S1G). Genera including Fusobacterium, Prevotella, and Leptotrichia had higher degree and betweenness centrality in CP-NA networks. These are correlation-network features rather than evidence of direct microbial interactions.

The CP-AD network showed a distinct topology, with lower modularity than the PH-NA network and a greater concentration of centrality in specific taxa. Anaerobic genera including Porphyromonas, Tannerella, and Treponema had high centrality measures. Differences in network properties describe niche-associated variation in correlation structure; they do not establish ecological rewiring or causation (Supplementary Table S1G).

### 5. Differentially abundant genera across periodontal niches

Differential abundance analysis identified multiple genera exhibiting consistent differences across periodontal niches (Fig. 3D–F). Compared with PH-NA samples, CP-NA samples showed significantly reduced relative abundances of commensal genera such as Gemella, Neisseria, and Lautropia. In parallel, CP-NA samples exhibited increased relative abundance of Fusobacterium, which was consistently detected across individuals. Comparison between PH-NA and CP-AD samples revealed broader taxonomic differences. CP-AD samples were enriched in obligate anaerobic genera, including Porphyromonas, Tannerella, Treponema, Fretibacterium, and Desulfobulbus, whereas health-associated genera were markedly depleted. These lesion-associated genera were among the most consistently enriched taxa across CP-AD samples.

Direct comparison between CP-NA and CP-AD highlighted niche-associated differences within periodontitis plaque. Genera such as Porphyromonas, Tannerella, and Treponema had higher relative abundance in CP-AD samples, whereas Leptotrichia and Campylobacter were relatively more abundant in CP-NA samples. Predictive modelling was not included because the archived model was fitted at the ASV level and subsequently mapped to genus labels; a genus-level, nested, patient-grouped validation would be required before reporting predictive performance.

## Discussion

We characterized subgingival microbial communities across distinct sampling niches using paired non-attached and attached plaque from participants with periodontitis, with healthy non-attached plaque as a reference. The analyses identified niche-associated differences in composition, diversity, and correlation-network structure. Across several analyses, CP-NA samples occupied positions between PH-NA and CP-AD samples. Because the study is cross-sectional, this ordering should be interpreted as a description of sampled niches rather than evidence of disease progression.

Previous studies have often contrasted subgingival microbiota between periodontal health and disease while treating subgingival plaque as a homogeneous entity [8,9]. The subgingival environment is spatially structured, with gradients in oxygen availability, shear forces, and proximity to host tissues. Non-attached plaque occupies a distinct physical position within the periodontal pocket [16,17]. Our findings indicate that this niche has a mixed composition, containing commensal genera commonly associated with health and anaerobic taxa commonly enriched in periodontitis.

Beyond taxonomic composition, correlation-network topology differed across periodontal niches. Health-associated plaque showed a more modular structure, whereas the attached-plaque network showed lower modularity and a higher maximum node-level betweenness value. The non-attached periodontitis network contained more retained associations and had higher density and mean degree than the healthy network. These analyses describe co-occurrence patterns and community organization; they do not infer direct microbial interactions or causal mechanisms.

The study has several limitations. Its cross-sectional design precludes inference about temporal dynamics or disease progression. The healthy comparison group was smaller than the periodontitis group, and the available clinical dataset did not support stage- or grade-specific analysis. Amplicon sequencing constrained taxonomic resolution and did not measure functional potential. Network analyses were based on correlation patterns and should be interpreted as descriptive. Future longitudinal studies integrating clinical severity measures, shotgun metagenomics, and host-derived data are needed to test the functional and temporal implications of these niche-associated patterns. Together, these findings provide a niche-resolved description of periodontal dysbiosis and generate hypotheses for longitudinal and mechanistic studies.

## Conclusions

Subgingival microbial communities in periodontitis showed niche-specific organization. Non-attached plaque from participants with periodontitis had a mixed microbial profile that differed from both healthy non-attached plaque and attached plaque. These cross-sectional findings support a niche-resolved view of periodontal dysbiosis but do not establish disease progression, microbial causality, or clinical utility.

## Supporting information

Supplementary_File_S2_Reproducibility

periodontitis_reproducibility_PRJNA1512026

## Data Availability

Raw 16S rRNA gene amplicon sequencing data (V3–V4 region) are being submitted to the NCBI Sequence Read Archive under BioProject accession PRJNA1512026 (16S rRNA gene sequencing of the subgingival microbiome in periodontitis). The BioProject and 82 BioSample records have been processed. Individual BioSample and SRA run accessions will be added when SRA processing is complete and the records are released. The analysis code, de-identified processed inputs, and statistical outputs are supplied in Supplementary File S2.

### Acknowledgments

The authors thank all participants for their involvement in this study and acknowledge the staff of the Department of Stomatology, Xiangya Second Hospital of Central South University, for their assistance with participant recruitment and sample collection. We also thank BGI Genomics for technical support in sequencing.

## Author Contributions

Qingxiu Li and Guangmei Li contributed equally to this work. Qingxiu Li contributed to study conceptualization, data analysis, interpretation, and manuscript drafting. Guangmei Li contributed to participant recruitment, clinical coordination, sample collection, and data curation. Zhenjun Li and Zhiwen Liu contributed to study design, supervision, and critical manuscript revision, and served as corresponding authors. All authors reviewed and approved the final manuscript.

