## Supplementary_Table_S1 for "Niche-specific microbial community structure of subgingival plaque in periodontitis"

### Supplementary Table S1A. Study design and sample structure

| Group | Sampling niche | Participants | Samples | Design note |
|---|---|---|---|---|
| PH-NA | Periodontally healthy non-attached plaque | 10 | 10 | Unpaired reference group |
| CP-NA | Periodontitis non-attached plaque | 36 | 36 | One of a paired sample set |
| CP-AD | Periodontitis attached plaque | 36 | 36 | Paired with CP-NA from the same participant |

CP-NA and CP-AD were collected as paired samples from the same 36 participants with periodontitis.

### Supplementary Table S1B. Shannon diversity summary

| Comparison set | Group | Samples | Mean Shannon | SD | Median | IQR | Minimum | Maximum |
|---|---|---|---|---|---|---|---|---|
| Healthy vs Periodontitis | Healthy | 10 | 3.2992 | 0.6818 | 3.1415 | 0.6414 | 2.3091 | 4.723 |
| Healthy vs Periodontitis | Periodontitis | 72 | 5.4318 | 0.3884 | 5.5178 | 0.5889 | 4.4249 | 6.0665 |
| CP-NA vs CP-AD | CP-NA | 36 | 5.2601 | 0.3401 | 5.262 | 0.5072 | 4.4566 | 5.9161 |
| CP-NA vs CP-AD | CP-AD | 36 | 5.6036 | 0.3601 | 5.6835 | 0.3435 | 4.4249 | 6.0665 |

Values are calculated from the archived analysis output. Each row reports sample-level Shannon diversity.

### Supplementary Table S1C. Design-aware Shannon diversity tests

| Comparison | Design | n (first group) | n (second group) | Mean Shannon (first group) | Mean Shannon (second group) | P value |
|---|---|---|---|---|---|---|
| PH-NA vs CP-NA | independent | 10 | 36 | 3.299178809931111 | 5.260056034908343 | 2.596477621366113e-06 |
| CP-NA vs CP-AD | paired | 36 | 36 | 5.260056034908343 | 5.603562342723986 | 2.566695911809802e-06 |

PH-NA versus CP-NA was analysed as an independent comparison; CP-NA versus CP-AD used a paired Wilcoxon signed-rank test.

### Supplementary Table S1D. Design-aware Bray–Curtis PERMANOVA and dispersion tests

| Comparison | Design | n (first group) | n (second group) | PERMANOVA F | PERMANOVA $R^2$ | PERMANOVA P | Dispersion test | Dispersion P |
|---|---|---|---|---|---|---|---|---|
| PH-NA vs CP-NA | independent | 10 | 36 | 23.3303 | 0.3465 | 0.001 | Mann–Whitney U | 0.0198 |
| CP-NA vs CP-AD | paired, labels permuted within participant | 36 | 36 | 6.1685 | 0.081 | 0.001 | Wilcoxon signed-rank | 0.0004 |

For CP-NA versus CP-AD, PERMANOVA labels were permuted only within participants. Significant dispersion tests require cautious interpretation of PERMANOVA as an overall distributional difference.

### Supplementary Table S1E. Chao1 richness summary

| Index | Group | Comparison set | Samples | Mean Chao1 | SD | Median | IQR | Minimum | Maximum |
|---|---|---|---|---|---|---|---|---|---|
| Chao1 | Healthy | Healthy vs Periodontitis | 10 | 257.8 | 116.3842 | 225.5 | 103.5 | 139.0 | 542.0 |
| Chao1 | Periodontitis | Healthy vs Periodontitis | 72 | 421.8801 | 103.1941 | 432.7308 | 131.5395 | 153.1 | 634.1667 |
| Chao1 | CP-NA | CP-NA vs CP-AD | 36 | 373.329 | 92.7045 | 390.375 | 118.6979 | 153.1 | 552.2069 |
| Chao1 | CP-AD | CP-NA vs CP-AD | 36 | 470.4311 | 90.3275 | 474.0 | 121.7496 | 254.0 | 634.1667 |

Chao1 richness was calculated from the ASV count matrix. Values are reported for the same samples as the Shannon summaries.

### Supplementary Table S1F. Design-aware Chao1 richness tests

| Comparison | Design | n (first group) | n (second group) | Mean Chao1 (first group) | Mean Chao1 (second group) | P value |
|---|---|---|---|---|---|---|
| PH-NA vs CP-NA | independent | 10 | 36 | 257.8 | 373.3290342923107 | 0.0032531560965154 |
| CP-NA vs CP-AD | paired | 36 | 36 | 373.3290342923107 | 470.4310736028641 | 8.043716661632061e-07 |

PH-NA versus CP-NA was analysed as an independent comparison; CP-NA versus CP-AD used a paired Wilcoxon signed-rank test. P values are unadjusted, two-sided values for prespecified comparisons.

### Supplementary Table S1G. Genus-level association-network topology

| Group | Nodes | Retained edges | Density | Mean degree | Louvain modularity | Maximum normalized | Positive edges | Negative edges |
|---|---|---|---|---|---|---|---|---|

| | | | | | | **betweenness** | | |
|---|---|---|---|---|---|---|---|---|
| PH-NA | 22 | 24 | 0.1039 | 2.1818 | 0.7431 | 0.019 | 24 | 0 |
| CP-NA | 54 | 180 | 0.1258 | 6.6667 | 0.2834 | 0.1009 | 120 | 60 |
| CP-AD | 54 | 125 | 0.0874 | 4.6296 | 0.4428 | 0.2433 | 91 | 34 |

Networks were calculated within each group using genera present in at least 20% of group samples. Retained edges met |ρ| ≥ 0.5 and Benjamini–Hochberg-adjusted $P < 0.05$. Modularity uses the Louvain partition; maximum normalized betweenness summarizes the greatest node-level betweenness in each network.
