## Supplementary_File_S2_Reproducibility for "Niche-specific microbial community structure of subgingival plaque in periodontitis"

### Supplementary File S2. Reproducibility materials

#### Contents

The accompanying archive, `periodontitis_reproducibility_PRJNA1512026.zip`, contains the following de-identified analysis inputs, scripts, and outputs:

- `inputs/metadata.xlsx` and `inputs/ASV_taxonomy_table.csv`: de-identified analysis metadata and ASV taxonomy/count inputs.
- `code/paired_design_reanalysis.py`: design-aware Shannon/Chao1 diversity and Bray–Curtis PERMANOVA analyses. PH-NA versus CP-NA is treated as independent; CP-NA versus CP-AD is paired by participant.
- `outputs/paired_design_alpha_diversity.tsv`, `outputs/paired_design_chao1_diversity.tsv`, and `outputs/paired_design_beta_diversity.tsv`: tabulated design-aware statistical results.
- `code/genus_association_network.R` and `outputs/net_edges_*.csv`: genus-level Spearman association-network code and retained edge tables. Genera present in at least 20% of samples within a group were retained; network edges met |rho| ≥ 0.5 and Benjamini–Hochberg-adjusted $P < 0.05$.
- `code/forest_plot.R`: genus-level relative-abundance comparisons, Hodges–Lehmann effect sizes, confidence intervals, and Benjamini–Hochberg adjustment used for Figure 3D–F.
- `code/phylogenetic_tree.R`, `code/genus_heatmap.R`, and `code/venn_diagram.R`: scripts used for the phylogenetic and taxonomic-composition visualizations.

#### Software

Primary sequence processing used DADA2 v1.28.0 in R v4.3.2 and SILVA release 138.1. The downstream R scripts state their package imports and random seeds. The paired-design Python analysis requires Python with NumPy, pandas, SciPy, and openpyxl. The archive includes `README.md` describing its contents and limits.

#### Data access

Raw 16S rRNA V3–V4 reads are being submitted to the NCBI Sequence Read Archive under BioProject PRJNA1512026. The BioProject and 82 BioSample records have been processed. Individual SRA run and BioSample accessions will be added when SRA processing and release are complete.
