## Supplementary material for "Niche-specific microbial community structure of subgingival plaque in periodontitis": periodontitis_reproducibility_PRJNA1512026: dada2_asv_processing.R

```
library(dada2)
library(Biostrings)

# 设置路径
path <- "C:/Users/84329/Desktop/宏基因组/牙周炎/data/fq"
filt_path <- file.path(path, "filtered_v3v4_final")
if (!dir.exists(filt_path)) dir.create(filt_path)

# 获取所有前向文件
fnFs <- sort(list.files(path, pattern = "_1.fq$", full.names = TRUE))
fnRs <- sort(list.files(path, pattern = "_2.fq$", full.names = TRUE))

# 提取样本名（去掉_1.fq）
sample.names <- gsub("_1.fq", "", basename(fnFs))


# 输出文件路径
filtFs <- file.path(filt_path, paste0(sample.names, "_F_filt.fastq.gz"))
filtRs <- file.path(filt_path, paste0(sample.names, "_R_filt.fastq.gz"))

# 执行过滤（推荐 truncLen 适用于 V3–V4）
out <- filterAndTrim(fnFs, filtFs, fnRs, filtRs,
                     truncLen = c(260, 220),
                     maxN = 0, maxEE = c(2, 2), truncQ = 2,
                     rm.phix = TRUE, compress = TRUE, multithread = TRUE)
print(out)

errF <- learnErrors(filtFs, multithread = TRUE)
errR <- learnErrors(filtRs, multithread = TRUE)
names(filtFs) <- sample.names
names(filtRs) <- sample.names

dadaFs <- vector("list", length(sample.names))
dadaRs <- vector("list", length(sample.names))
mergers <- vector("list", length(sample.names))

for (i in seq_along(sample.names)) {
  derepF <- derepFastq(filtFs[i])
  derepR <- derepFastq(filtRs[i])
  dadaFs[[i]] <- dada(derepF, err = errF, multithread = FALSE)
  dadaRs[[i]] <- dada(derepR, err = errR, multithread = FALSE)
  mergers[[i]] <- mergePairs(dadaFs[[i]], derepF, dadaRs[[i]], derepR)
}

names(mergers) <- sample.names

# 构建全表
seqtab <- makeSequenceTable(mergers)

# 去除低丰度ASV（如总丰度 ≤10）
asv_sums <- colSums(seqtab)
seqtab.filt <- seqtab[, asv_sums > 10]

# 去除嵌合体
seqtab.nochim <- removeBimeraDenovo(seqtab.filt, method = "consensus", multithread = TRUE)

# 保存
saveRDS(seqtab.nochim, file.path(path, "seqtab_nochim.rds"))

ref_fasta <- file.path(path, "eHOMD_RefSeq_dada2_V15.22.fasta")
taxa <- assignTaxonomy(seqtab.nochim, refFasta = ref_fasta, multithread = TRUE)

# 保存注释结果
saveRDS(taxa, file.path(path, "taxonomy.rds"))

asv_seqs <- colnames(seqtab.nochim)
asv_tab <- t(seqtab.nochim)
asv_table <- data.frame(ASV = asv_seqs, asv_tab, taxa)
write.csv(asv_table, file.path(path, "ASV_taxonomy_table.csv"), row.names = FALSE)
```
