## Supplementary material for "Niche-specific microbial community structure of subgingival plaque in periodontitis": periodontitis_reproducibility_PRJNA1512026: forest_plot.R

```
## =========================================================
## 三幅森林图：PH-NA vs CP-NA（非配对）、PH-NA vs CP-AD（非配对）、CP-NA vs CP-AD（配对）
## 输入：metadata.xlsx（需列 Number, Sampling_Location；配对比较还需 PatientID）
##       ASV_taxonomy_table.csv（Genus + 样本列）
## 输出：Perio_forest_HL_topN_PH-NA_vs_CP-NA_NC.{png,pdf}
##       Perio_forest_HL_topN_PH-NA_vs_CP-AD_NC.{png,pdf}
##       Perio_forest_HL_topN_CP-NA_vs_CP-AD_NC.{png,pdf}
## =========================================================
suppressPackageStartupMessages({
  library(readxl)
  library(dplyr); library(tidyr); library(stringr)
  library(ggplot2); library(readr)
})

## ---------------- 路径 & 参数 ----------------
meta_file <- "metadata.xlsx"
asv_file  <- "ASV_taxonomy_table.csv"

TopN <- 25
grp_levels <- c("PH-NA","CP-NA","CP-AD")
cols_grp   <- c("PH-NA"="#009E73","CP-NA"="#0072B2","CP-AD"="#CC79A7")

## ---------------- 辅助函数 ----------------
clean_group <- function(x){
  y <- x %>% stringr::str_trim() %>% stringr::str_replace_all("\\s+","") %>%
    toupper() %>% stringr::str_replace_all("_","-")
  dplyr::case_when(
    y %in% c("PH-NA","PHNA","H-NA","HNA","HEALTHY","HEALTHYNA","PH") ~ "PH-NA",
    y %in% c("CP-NA","CPNA","NONATTACHED","NASP","N-SUB","NSUB")      ~ "CP-NA",
    y %in% c("CP-AD","CPAD","ATTACHED","ASP","A-SUB","ASUB")          ~ "CP-AD",
    TRUE ~ NA_character_
  )
}

safe_div <- function(m, totals){        # 列归一化（相对丰度 %）
  out <- m; z <- totals == 0 | is.na(totals)
  if (any(!z)) out[, !z] <- sweep(m[, !z, drop = FALSE], 2, totals[!z], "/")
  if (any(z))  out[,  z] <- 0
  out * 100
}

clean_shorten <- function(x, k = 12){   # 属名缩写
  y <- gsub("\\[(.*?)\\]", "\\1", x)
  y <- gsub("[\\(\\)\\[\\]_\\-\\s]+", "", y)
  y <- stringr::str_sub(y, 1, k)
  make.unique(y, sep=".")
}

## HL（非配对：所有 pairwise 差的中位数）
HL_unpaired <- function(x, y){
  if (length(x) == 0 || length(y) == 0) return(NA_real_)
  median(outer(x, y, `-`), na.rm = TRUE)
}

## 简单 bootstrap CI 兜底
boot_ci <- function(x, y = NULL, paired = FALSE, B = 2000, conf = 0.95){
  if (paired) {
    stopifnot(length(x) == length(y))
    d <- x - y
    n <- length(d)
    if (n < 3) return(c(NA_real_, NA_real_))
    idx <- matrix(sample.int(n, n * B, replace = TRUE), nrow = n)
    stats <- apply(idx, 2, function(i) median(d[i], na.rm = TRUE))
  } else {
    if (length(x) < 2 || length(y) < 2) return(c(NA_real_, NA_real_))
    nx <- length(x); ny <- length(y)
    stats <- replicate(B, {
      xb <- sample(x, nx, replace = TRUE)
      yb <- sample(y, ny, replace = TRUE)
      HL_unpaired(xb, yb)
    })
  }
  alpha <- (1 - conf)/2
  stats <- stats[is.finite(stats)]
  if (length(stats) < 10) return(c(NA_real_, NA_real_))
  quantile(stats, probs = c(alpha, 1 - alpha), names = FALSE, type = 6)
}

## 统一返回 hl/lwr/upr/p；paired=TRUE 用配对 Wilcoxon
wilcox_hl_adapt <- function(a, b, paired = FALSE){
  if (paired) {
    if (length(a) == 0 || length(b) == 0) return(c(hl=NA, lwr=NA, upr=NA, p=NA))
    wt <- tryCatch(
      stats::wilcox.test(a, b, paired = TRUE, conf.int = TRUE, exact = FALSE),
      error = function(e) NULL
    )
    if (is.null(wt)) return(c(hl=NA, lwr=NA, upr=NA, p=NA))
    hl  <- unname(if (!is.null(wt$estimate)) wt$estimate else median(a - b, na.rm = TRUE))
    ci  <- if (!is.null(wt$conf.int)) unname(wt$conf.int) else boot_ci(a, b, paired = TRUE)
    p   <- wt$p.value
    c(hl = hl, lwr = ci[1], upr = ci[2], p = p)
  } else {
    hl <- HL_unpaired(a, b)
    wt <- tryCatch(
      stats::wilcox.test(a, b, conf.int = TRUE, exact = FALSE),
      error = function(e) NULL
    )
    if (!is.null(wt) && !is.null(wt$conf.int)) {
      ci <- unname(wt$conf.int)
      p  <- wt$p.value
    } else {
      ci <- boot_ci(a, b, paired = FALSE)
      p  <- if (!is.null(wt)) wt$p.value else NA_real_
    }
    c(hl = hl, lwr = ci[1], upr = ci[2], p = p)
  }
}

## ---------------- 读取 metadata ----------------
meta_raw <- readxl::read_xlsx(meta_file)
if (!all(c("Number","Sampling_Location") %in% names(meta_raw))) {
  stop("metadata.xlsx 必须包含列 Number 与 Sampling_Location")
}
has_pid <- "PatientID" %in% names(meta_raw)

meta <- meta_raw %>%
  mutate(
    Number = as.character(.data$Number),
    Group  = clean_group(.data$Sampling_Location),
    PatientID = if (has_pid) as.character(.data$PatientID) else NA_character_
  ) %>%
  filter(!is.na(.data$Number), !is.na(.data$Group)) %>%
  filter(.data$Group %in% grp_levels) %>%
  mutate(Group = factor(.data$Group, levels = grp_levels)) %>%
  select(.data$Number, .data$Group, .data$PatientID)

## 若没有 PatientID，则无法做配对比较
if (!has_pid || all(is.na(meta$PatientID))) {
  stop("CP-NA vs CP-AD 需要 PatientID 进行配对检验：请在 metadata.xlsx 添加 PatientID 列。")
}

## ---------------- 读取 ASV 表并聚合到 Genus 相对丰度 ----------------
asv_tab <- utils::read.csv(asv_file, check.names = FALSE, stringsAsFactors = FALSE)
if (!("Genus" %in% names(asv_tab))) stop("ASV_taxonomy_table.csv 必须含列 'Genus'。")

sample_cols <- intersect(colnames(asv_tab), meta$Number)
if (length(sample_cols) == 0) stop("ASV 表未找到与 metadata$Number 匹配的样本列。")

asv_tab$Genus <- as.character(asv_tab$Genus)
asv_tab$Genus[is.na(asv_tab$Genus) | asv_tab$Genus==""] <- "Unassigned"

for (cn in sample_cols) {
  v <- asv_tab[[cn]]
  asv_tab[[cn]] <- suppressWarnings(as.numeric(v))
  asv_tab[[cn]][is.na(asv_tab[[cn]])] <- 0
}

genus_mat <- asv_tab %>%
  select(.data$Genus, dplyr::all_of(sample_cols)) %>%
  group_by(.data$Genus) %>%
  summarise(across(all_of(sample_cols), ~ sum(.x, na.rm = TRUE)), .groups = "drop")

totals <- colSums(as.matrix(genus_mat[, sample_cols, drop = FALSE]), na.rm = TRUE)
genus_rel <- genus_mat
genus_rel[, sample_cols] <- safe_div(as.matrix(genus_mat[, sample_cols, drop = FALSE]), totals)

long_df <- genus_rel %>%
  tidyr::pivot_longer(cols = dplyr::all_of(sample_cols),
                      names_to = "Number", values_to = "RelAbund") %>%
  left_join(meta, by = "Number") %>%
  filter(!is.na(.data$Group))

## ---------------- 三个比较的定义与配对模式 ----------------
cmp_list <- list(
  `PH-NA vs CP-NA` = list(L="PH-NA", R="CP-NA", paired=FALSE),
  `PH-NA vs CP-AD` = list(L="PH-NA", R="CP-AD", paired=FALSE),
  `CP-NA vs CP-AD` = list(L="CP-NA", R="CP-AD", paired=TRUE )
)

## ---------------- 单个比较计算 & 绘图函数 ----------------
run_one_comparison <- function(L, R, paired_flag, topN = TopN){
  sub <- long_df %>% filter(Group %in% c(L, R))
  gens <- unique(sub$Genus)
  
  res_list <- lapply(gens, function(g){
    gdat <- sub %>% filter(Genus == g)
    
    if (paired_flag) {
      wide <- gdat %>%
        select(PatientID, Group, RelAbund) %>%
        filter(!is.na(PatientID)) %>%
        tidyr::pivot_wider(names_from = Group, values_from = RelAbund)
      wide <- wide %>% filter(!is.na(.data[[L]]), !is.na(.data[[R]]))
      vL <- wide[[L]]; vR <- wide[[R]]
      kv <- wilcox_hl_adapt(vR, vL, paired = TRUE)   # HL = Right − Left
    } else {
      vL <- gdat$RelAbund[gdat$Group == L]
      vR <- gdat$RelAbund[gdat$Group == R]
      kv <- wilcox_hl_adapt(vR, vL, paired = FALSE)  # HL = Right − Left
    }
    
    data.frame(
      Genus = g,
      hl  = kv["hl"], lwr = kv["lwr"], upr = kv["upr"], p = kv["p"],
      Left = L, Right = R,
      stringsAsFactors = FALSE
    )
  })
  
  df <- bind_rows(res_list)
  df$FDR <- stats::p.adjust(df$p, method = "BH")
  df$Comparison <- paste(L, "vs", R)
  df$winner <- ifelse(df$hl >= 0, df$Right, df$Left)
  df$winner <- factor(df$winner, levels = grp_levels)
  
  ## TopN & 排序（仅在该比较内部）
  df2 <- df %>%
    arrange(FDR, desc(abs(hl))) %>%
    mutate(Genus_short = clean_shorten(.data$Genus, k = 12)) %>%
    slice_head(n = topN)
  
  ## 为本比较单独生成 y 因子水平
  ord <- df2 %>% arrange(FDR, desc(abs(hl))) %>% pull(Genus_short) %>% unique()
  df2$y_fac <- factor(df2$Genus_short, levels = rev(ord))
  df2
}

plot_one_comparison <- function(df_cmp){
  cmp <- unique(df_cmp$Comparison)
  stopifnot(length(cmp) == 1)
  L <- unique(df_cmp$Left);  R <- unique(df_cmp$Right)
  
  p <- ggplot(df_cmp, aes(x = .data$hl, y = .data$y_fac, color = .data$winner)) +
    geom_vline(xintercept = 0, linewidth = 0.3, linetype = 2) +
    geom_errorbarh(aes(xmin = .data$lwr, xmax = .data$upr), height = 0, linewidth = 0.7, alpha = 0.7) +
    geom_point(size = 2) +
    scale_color_manual(values = cols_grp, name = "Winner group") +
    labs(
      x = sprintf("Hodges–Lehmann effect (Δ%%, %s Right − Left)", cmp),
      y = NULL,
      title = paste0("Top differential genera — ", cmp)
    ) +
    theme_classic(base_size = 10) +
    theme(
      axis.line.y = element_blank(),
      legend.position = "right",
      plot.title = element_text(hjust = 0.5, face = "bold")
    )
  p
}

## ---------------- 逐一运行并出图 ----------------
all_outputs <- list()
for (nm in names(cmp_list)) {
  L <- cmp_list[[nm]]$L; R <- cmp_list[[nm]]$R; paired_flag <- cmp_list[[nm]]$paired
  message("Computing: ", nm, " ...")
  df_cmp <- run_one_comparison(L, R, paired_flag, topN = TopN)
  all_outputs[[nm]] <- df_cmp
  
  p <- plot_one_comparison(df_cmp)
  
  file_stub <- gsub("\\s+","", gsub("[^A-Za-z0-9]+","_", nm))
  ggsave(paste0("Perio_forest_HL_topN_", file_stub, "_NC.png"),
         p, width = 160, height = 130, units = "mm", dpi = 600, bg = "white")
  ggsave(paste0("Perio_forest_HL_topN_", file_stub, "_NC.pdf"),
         p, width = 160, height = 130, units = "mm", device = cairo_pdf, bg = "transparent")
}

## 可选：合并结果保存一份 CSV
bind_rows(all_outputs) %>% write_csv("functional_candidates_perio.csv")

message("OK：已按要求生成 3 幅森林图（两幅非配对 + 一幅配对），并导出 CSV。")
```
