## Supplementary material for "Niche-specific microbial community structure of subgingival plaque in periodontitis": periodontitis_reproducibility_PRJNA1512026: genus_association_network.R

```
## =========================================================
## 三组（PH-NA / CP-NA / CP-AD）—— Genus 相关性网络（Base R，无 dplyr/tidyr）
## 依赖：readxl, Hmisc, igraph, ggraph, ggplot2
## 输入：metadata.xlsx（Number, Sampling_Location）、ASV_taxonomy_table.csv（Genus + 样本列）
## 输出：net_edges_<GROUP>.csv / net_nodes_<GROUP>.csv /
##      network_genus_<GROUP>_{FR|RING}_NC.(png|pdf|svg)
## =========================================================

suppressPackageStartupMessages({
  library(readxl)
  library(Hmisc)    # rcorr() Spearman
  library(igraph)   # 图模型
  library(ggraph)   # 网络绘图
  library(ggplot2)
})

options(stringsAsFactors = FALSE)

## ------------ 参数 ------------
meta_file <- "metadata.xlsx"
asv_file  <- "ASV_taxonomy_table.csv"

## 组别与配色（固定顺序：健康在前）
groups_all <- c("PH-NA","CP-NA","CP-AD")

## 过滤与阈值
prev_thr   <- 0.20   # 出现率阈值（组内 >0 样本占比 ≥ 20%）
rho_thr    <- 0.5    # |rho| 阈值
fdr_thr    <- 0.05   # FDR 阈值
min_samples_for_corr <- 6  # 相关性至少需要的样本数（建议≥6）

## 图形参数
only_show_connected <- TRUE
col_edge <- c(pos = "#D55E00", neg = "#009E73")               # 边：红=正、绿=负
node_pal <- c(low="#E0E0E0", high="#6A3D9A")                  # 点：prevalence 连续色
fig_w_mm <- 220; fig_h_mm <- 170; dpi_out <- 600

## ------------ 小工具 ------------
clean_group <- function(x){
  y <- toupper(gsub("\\s+","", gsub("_","-", trimws(as.character(x)))))
  y[y %in% c("PH-NA","PHNA","H-NA","HNA","HEALTHY","HEALTHYNA","PH")] <- "PH-NA"
  y[y %in% c("CP-NA","CPNA","NONATTACHED","NASP","N-SUB","NSUB")]     <- "CP-NA"
  y[y %in% c("CP-AD","CPAD","ATTACHED","ASP","A-SUB","ASUB")]         <- "CP-AD"
  y
}
shorten_label <- function(x, k = 6){
  y <- gsub("\\[(.*?)\\]", "\\1", x)
  y <- gsub("[\\(\\)\\[\\]_\\-\\s]+", "", y)
  y <- substr(y, 1, k)
  make.unique(y, sep=".")
}
rescale_vec <- function(x, to=c(3,9)){
  rng <- range(x, na.rm=TRUE)
  if (!is.finite(rng[1]) || rng[1]==rng[2]) return(rep(mean(to), length(x)))
  (x - rng[1])/(rng[2]-rng[1]) * diff(to) + to[1]
}

## =========================================================
## 读取 metadata & ASV 表（一次）
## =========================================================
meta <- read_xlsx(meta_file)
if (!all(c("Number","Sampling_Location") %in% names(meta)))
  stop("metadata.xlsx 必须包含列：Number, Sampling_Location")

meta$Number <- as.character(meta$Number)
meta$Group  <- clean_group(meta$Sampling_Location)
meta <- meta[meta$Group %in% groups_all & !is.na(meta$Number), c("Number","Group")]
if (!nrow(meta)) stop("metadata 中无 PH-NA/CP-NA/CP-AD 可用记录。")

asv <- read.csv(asv_file, check.names = FALSE)
if (!("Genus" %in% names(asv))) stop("ASV 表缺少列：Genus")
asv$Genus <- as.character(asv$Genus)
asv <- asv[!is.na(asv$Genus) & nzchar(asv$Genus), , drop=FALSE]

## 找所有样本列（与 meta$Number 交集）
sample_cols_all <- intersect(colnames(asv), unique(meta$Number))
if (!length(sample_cols_all)) stop("ASV 表中没有与 metadata$Number 匹配的样本列。")

## 确保样本列为数值
for (cn in sample_cols_all) asv[[cn]] <- suppressWarnings(as.numeric(asv[[cn]]))

## =========================================================
## 核心函数：构建“组内”Genus 相关性网络
## =========================================================
build_group_network <- function(GRP){
  message(">>> 处理组：", GRP)
  smps <- meta$Number[meta$Group == GRP]
  smps <- intersect(smps, sample_cols_all)
  if (length(smps) < min_samples_for_corr) {
    warning(GRP, ": 样本数 < ", min_samples_for_corr, "；跳过该组网络。")
    return(invisible(NULL))
  }
  
  ## 1) 聚合到 Genus（Base R aggregate）
  keep <- c("Genus", smps)
  tmp  <- asv[, keep, drop=FALSE]
  genus_mat <- aggregate(x = tmp[, smps, drop=FALSE],
                         by = list(Genus = tmp$Genus),
                         FUN = function(z) sum(z, na.rm=TRUE))
  
  ## 2) 转相对丰度（%）
  cs <- colSums(genus_mat[, smps, drop=FALSE], na.rm=TRUE); cs[cs==0] <- NA_real_
  genus_rel <- genus_mat
  genus_rel[, smps] <- sweep(genus_mat[, smps, drop=FALSE], 2, cs, "/") * 100
  
  ## 3) 过滤低出现率属（组内 >0 的样本占比）
  M <- as.matrix(genus_rel[, smps, drop=FALSE]); rownames(M) <- genus_rel$Genus
  prev <- rowMeans(M > 0, na.rm=TRUE)
  keep_rows <- which(prev >= prev_thr)
  if (!length(keep_rows)) {
    warning(GRP, ": 过滤后无属可用于相关分析。")
    return(invisible(NULL))
  }
  M <- M[keep_rows, , drop=FALSE]
  
  ## 4) 相关性（变量=属 ⇒ 列；样本为观测 ⇒ 行）
  X <- t(M)  # 行=样本, 列=Genus
  rc <- Hmisc::rcorr(X, type = "spearman")
  r_mat <- rc$r; p_mat <- rc$P
  
  ## 5) 上三角展开 + FDR + 阈值
  ut <- which(upper.tri(r_mat), arr.ind = TRUE)
  edges <- data.frame(
    Genus1 = colnames(r_mat)[ut[,1]],
    Genus2 = colnames(r_mat)[ut[,2]],
    rho    = r_mat[ut],
    pval   = p_mat[ut],
    stringsAsFactors = FALSE
  )
  edges <- edges[!is.na(edges$rho) & !is.na(edges$pval), , drop=FALSE]
  if (!nrow(edges)) {
    warning(GRP, ": 相关性结果全 NA；跳过绘图。")
    return(invisible(NULL))
  }
  edges$padj <- p.adjust(edges$pval, method = "BH")
  edges <- edges[abs(edges$rho) >= rho_thr & edges$padj < fdr_thr, , drop=FALSE]
  if (!nrow(edges)) {
    warning(GRP, ": 无满足 |rho|≥", rho_thr, " 且 FDR<", fdr_thr, " 的边；跳过绘图。")
    return(invisible(NULL))
  }
  edges$edge_sign <- ifelse(edges$rho >= 0, "pos", "neg")
  rownames(edges) <- NULL
  
  ## 6) 节点表（prevalence / mean_abund / 缩放大小 / 标签）
  g_names <- unique(c(edges$Genus1, edges$Genus2))
  Msub <- M[g_names, , drop=FALSE]
  mean_abund <- rowMeans(Msub, na.rm=TRUE)
  prevalence <- rowMeans(Msub > 0, na.rm=TRUE)
  nodes <- data.frame(
    name       = names(mean_abund),
    mean_abund = as.numeric(mean_abund),
    prevalence = as.numeric(prevalence),
    stringsAsFactors = FALSE
  )
  nodes$size  <- rescale_vec(nodes$mean_abund, to = c(3, 10))
  nodes$label <- shorten_label(nodes$name, k = 6)
  
  ## 7) 保存 edges/nodes
  write.csv(edges, paste0("net_edges_", GRP, ".csv"), row.names = FALSE)
  write.csv(nodes, paste0("net_nodes_", GRP, ".csv"), row.names = FALSE)
  
  ## 8) 画图 —— FR 布局
  g <- graph_from_data_frame(edges, vertices = nodes, directed = FALSE)
  p_fr <- ggraph(g, layout = "fr") +
    ggraph::geom_edge_link(aes(width = abs(rho), colour = edge_sign), alpha = 0.8) +
    ggraph::scale_edge_width(name = "|ρ|", range = c(0.3, 2.8)) +
    ggraph::scale_edge_colour_manual(values = col_edge, name = "ρ sign") +
    ggraph::geom_node_point(aes(size = size, colour = prevalence)) +
    scale_size_identity(guide = "none") +
    scale_colour_gradient(low = node_pal[1], high = node_pal[2], name = "Prevalence") +
    ggraph::geom_node_text(aes(label = label), repel = TRUE, size = 3, lineheight = 0.95) +
    theme_void(base_size = 10) +
    labs(title = paste0("Genus correlation network — ", GRP)) +
    theme(plot.title = element_text(hjust = 0.5, face = "bold"),
          legend.position = "right")
  ggsave(paste0("network_genus_", GRP, "_FR_NC.png"), p_fr,
         width = fig_w_mm, height = fig_h_mm, units = "mm", dpi = dpi_out, bg = "white")
  ggsave(paste0("network_genus_", GRP, "_FR_NC.pdf"), p_fr,
         width = fig_w_mm, height = fig_h_mm, units = "mm", device = cairo_pdf, bg = "transparent")
  
  ## 9) 画图 —— 环形弧线布局
  ##    先按 prevalence/mean_abund/label 排序，让外圈标签更有序
  nodes_ord <- nodes[order(-nodes$prevalence, -nodes$mean_abund, nodes$label), , drop=FALSE]
  g_ring <- graph_from_data_frame(edges, vertices = nodes_ord, directed = FALSE)
  
  p_ring <- ggraph(g_ring, layout = "linear", circular = TRUE) +
    # 背景浅灰弧线
    ggraph::geom_edge_arc(aes(width = abs(rho)), colour = "grey85", alpha = 0.6) +
    # 主弧线（正红负绿）
    ggraph::geom_edge_arc(aes(width = abs(rho), colour = edge_sign), alpha = 0.95) +
    ggraph::scale_edge_width(name = "|ρ|", range = c(0.3, 2.8)) +
    ggraph::scale_edge_colour_manual(values = col_edge, name = "ρ sign") +
    # 节点
    ggraph::geom_node_point(aes(size = size, colour = prevalence)) +
    scale_size_identity(guide = "none") +
    scale_colour_gradient(low = node_pal[1], high = node_pal[2], name = "Prevalence") +
    # 外圈标签（自动翻转角度）
    ggraph::geom_node_text(
      data = function(df) {
        ang  <- atan2(df$y, df$x) * 180 / pi
        flip <- ang < -90 | ang > 90
        data.frame(df,
                   lx = df$x * 1.06,
                   ly = df$y * 1.06,
                   angle = ifelse(flip, ang + 180, ang),
                   hjust = ifelse(flip, 1, 0))
      },
      aes(x = lx, y = ly, label = label, angle = angle, hjust = hjust),
      size = 3, lineheight = 0.95
    ) +
    coord_fixed(clip = "off") +
    theme_void(base_size = 10) +
    labs(title = paste0("Genus correlation network (ring) — ", GRP)) +
    theme(plot.title = element_text(hjust = 0.5, face = "bold"),
          legend.position = "right",
          plot.margin = margin(12, 20, 12, 16))
  ggsave(paste0("network_genus_", GRP, "_RING_NC.png"), p_ring,
         width = fig_w_mm, height = fig_h_mm, units = "mm", dpi = dpi_out, bg = "white")
  ggsave(paste0("network_genus_", GRP, "_RING_NC.pdf"), p_ring,
         width = fig_w_mm, height = fig_h_mm, units = "mm", device = cairo_pdf, bg = "transparent")
  ggsave(paste0("network_genus_", GRP, "_RING_NC.svg"), p_ring,
         width = fig_w_mm, height = fig_h_mm, units = "mm", bg = "transparent")
  
  invisible(list(edges = edges, nodes = nodes))
}

## =========================================================
## 逐组生成网络
## =========================================================
for (grp in groups_all) {
  try(build_group_network(grp), silent = TRUE)
}

message("完成：三组分组内相关性网络（CSV + 图件）已导出。")
```
