## Supplementary material for "Niche-specific microbial community structure of subgingival plaque in periodontitis": periodontitis_reproducibility_PRJNA1512026: genus_heatmap.R

```
## =========================================================
## Genus 热图（每组独立渐变；组内聚类；列聚类树；Phylum 顶部条带；右侧整图图例）
## - 牙周炎三水平：PH-NA（健康龈下）、CP-NA（炎-非附着）、CP-AD（炎-附着）
## - 属名后缀去重（如 Saccha.5）；避免 NA
## - 门图例标签去掉 "_" 后部分；缺失门色自动补齐；Others 如出现固定最后
## - 三个色条置于第 3 图下；第 3 图属名旋转 45°
## =========================================================

suppressPackageStartupMessages({
  library(readxl); library(dplyr); library(tidyr); library(ggplot2); library(stringr)
  library(patchwork); library(cowplot); library(grid); library(ggdendro); library(dendextend)
})

## 可选：如已安装 conflicted，可优先 dplyr（避免 MASS/plyr 顶替）
## if (requireNamespace("conflicted", quietly = TRUE)) {
##   conflicted::conflict_prefer("select","dplyr")
##   conflicted::conflict_prefer("filter","dplyr")
##   conflicted::conflict_prefer("mutate","dplyr")
##   conflicted::conflict_prefer("summarise","dplyr")
##   conflicted::conflict_prefer("arrange","dplyr")
## }

## ============ 参数 ============
meta_file <- "metadata.xlsx"
asv_file  <- "ASV_taxonomy_table.csv"

topN_genus     <- 50
loc_levels     <- c("PH-NA","CP-NA","CP-AD")   # 固定顺序：健康在前
transform_mode <- "log10"
pseudo_count   <- 1e-4
clip_quantiles <- c(0.01, 0.99)
heat_limits    <- NULL

## 每组主色（用于各自渐变：白→主色）
group_main_colors <- c("PH-NA"="#009E73", "CP-NA"="#0072B2", "CP-AD"="#CC79A7")

fig_width_mm  <- 260
fig_height_mm <- 170  # 为顶部树留一点空间

## 初始 Phylum 颜色（缺失将自动补色）
phylum_colors_base <- c(
  "Absonditabacteria" = "#F39B7FFF",
  "Actinobacteria"    = "#E64B35FF",
  "Bacteroidetes"     = "#4DBBD5FF",
  "Cyanobacteria"     = "#00A087FF",
  "Firmicutes"        = "#3C5488FF",
  "Fusobacteria"      = "#8491B4FF",
  "Gracilibacteria"   = "#91D1C2FF",
  "Proteobacteria"    = "#0073C2FF",
  "Saccharibacteria"  = "#7E6148FF",
  "Spirochaetes"      = "#925E9FFF",
  "Synergistetes"     = "#FFD600FF",
  "Unknown"           = "grey70"
)
others_color <- c("Others" = "lightgrey")

## ============ metadata ============
meta_raw <- readxl::read_xlsx(meta_file)
names(meta_raw) <- trimws(names(meta_raw))

clean_group <- function(x){
  y <- x %>% stringr::str_trim() %>% stringr::str_replace_all("\\s+","") %>%
    toupper() %>% stringr::str_replace_all("_","-")
  dplyr::case_when(
    y %in% c("PH-NA","PHNA","H-NA","HNA","HEALTHY","HEALTHYNA","PH") ~ "PH-NA",
    y %in% c("CP-NA","CPNA","NONATTACHED","NASP","N-SUB","NSUB") ~ "CP-NA",
    y %in% c("CP-AD","CPAD","ATTACHED","ASP","A-SUB","ASUB") ~ "CP-AD",
    TRUE ~ NA_character_
  )
}

meta <- meta_raw %>%
  dplyr::mutate(
    Number = as.character(.data$Number),
    Sampling_Location = clean_group(.data$Sampling_Location)
  ) %>%
  dplyr::select(dplyr::all_of(c("Number","Sampling_Location"))) %>%
  dplyr::filter(!is.na(.data$Number), !is.na(.data$Sampling_Location)) %>%
  dplyr::filter(.data$Sampling_Location %in% loc_levels) %>%
  dplyr::mutate(Sampling_Location = factor(.data$Sampling_Location, levels = loc_levels))

stopifnot(nrow(meta) > 0)

## ============ 读取 ASV × taxonomy 表 ============
asv_tab_raw <- utils::read.csv(asv_file, check.names = FALSE, stringsAsFactors = FALSE)

tax_cols <- intersect(colnames(asv_tab_raw),
                      c("Kingdom","Phylum","Class","Order","Family","Genus","Species"))
sample_cols_all <- setdiff(colnames(asv_tab_raw), tax_cols)
sample_cols <- intersect(sample_cols_all, meta$Number)
if (length(sample_cols) == 0) stop("未在 ASV 表中找到与 metadata$Number 重叠的样本列。")

asv_tab <- asv_tab_raw[, unique(c("Genus","Phylum", sample_cols)), drop = FALSE]
asv_tab$Genus  <- as.character(asv_tab$Genus)
asv_tab$Phylum <- as.character(asv_tab$Phylum)

## 重要：确保 Genus 非空，防止后续 NA
asv_tab$Genus[is.na(asv_tab$Genus) | asv_tab$Genus==""] <- "Unassigned"

## 数值化样本列
for (cn in sample_cols) {
  v <- asv_tab[[cn]]
  asv_tab[[cn]] <- suppressWarnings(as.numeric(v))
  asv_tab[[cn]][is.na(asv_tab[[cn]])] <- 0
}

## ============ Genus → Phylum 映射：按总丰度最大 ============
genus_phylum_map <- asv_tab %>%
  tidyr::pivot_longer(cols = dplyr::all_of(sample_cols),
                      names_to = "Sample", values_to = "Abund") %>%
  dplyr::mutate(Phylum = dplyr::if_else(is.na(.data$Phylum) | .data$Phylum=="", NA_character_, .data$Phylum)) %>%
  dplyr::group_by(.data$Genus, .data$Phylum) %>%
  dplyr::summarise(total = sum(.data$Abund, na.rm = TRUE), .groups = "drop_last") %>%
  dplyr::summarise(Phylum = .data$Phylum[which.max(.data$total)], .groups = "drop") %>%
  dplyr::mutate(Phylum = dplyr::if_else(is.na(.data$Phylum) | .data$Phylum=="", "Unknown", .data$Phylum))

## ============ 聚合到 Genus & 相对丰度 ============
genus_mat <- asv_tab %>%
  dplyr::group_by(.data$Genus) %>%
  dplyr::summarise(dplyr::across(dplyr::all_of(sample_cols), ~ sum(.x, na.rm = TRUE)), .groups = "drop")

common_samples <- intersect(meta$Number, colnames(genus_mat))
if (length(common_samples) == 0) stop("对齐后无共同样本，请检查样本编号。")
genus_mat <- genus_mat %>% dplyr::select(dplyr::all_of(c("Genus", common_samples)))
meta <- meta %>% dplyr::filter(.data$Number %in% common_samples)

sample_totals <- colSums(as.matrix(genus_mat[, common_samples, drop = FALSE]), na.rm = TRUE)
safe_div <- function(m, totals){
  out <- m; z <- totals == 0 | is.na(totals)
  if (any(!z)) out[, !z] <- sweep(m[, !z, drop = FALSE], 2, totals[!z], "/")
  if (any(z))  out[,  z] <- 0
  out * 100
}
genus_rel <- genus_mat
genus_rel[, common_samples] <- safe_div(as.matrix(genus_mat[, common_samples, drop = FALSE]), sample_totals)

## ============ Top-N Genus ============
genus_rel <- as.data.frame(genus_rel)
genus_rel$mean_abund <- rowMeans(genus_rel[, common_samples, drop = FALSE], na.rm = TRUE)
ord <- order(genus_rel$mean_abund, decreasing = TRUE, na.last = TRUE)
keepN <- if (is.finite(topN_genus)) min(topN_genus, nrow(genus_rel)) else nrow(genus_rel)
top_genus_full <- genus_rel$Genus[ord][seq_len(keepN)]

genus_rel_top <- genus_rel %>%
  dplyr::filter(.data$Genus %in% top_genus_full) %>%
  dplyr::select(dplyr::all_of(c("Genus", common_samples))) %>%
  dplyr::left_join(genus_phylum_map, by = "Genus")

## ============ 属名显示：稳健清洗 + 唯一后缀 .2/.3 ============
shorten_genus <- function(x){
  x <- ifelse(is.na(x) | x=="", "Unassigned", x)
  x <- stringr::str_squish(x)
  x <- stringr::str_replace_all(x, "\u00A0", " ")
  x <- stringr::str_replace(x, "^_+.*$", "Unassigned")  # 以 '_' 开头的极端情形
  x <- stringr::str_replace(x, "_.*", "")               # 去掉第一个 '_' 及其后的部分
  x <- stringr::str_replace_all(x, "_", ".")            # 其余 '_' → '.'
  x <- substr(x, 1, 6)
  ifelse(x=="", "Unassigned", x)
}

label_map <- genus_rel_top %>%
  dplyr::mutate(Genus = ifelse(is.na(.data$Genus) | .data$Genus=="", "Unassigned", .data$Genus)) %>%
  dplyr::distinct(.data$Genus, .data$Phylum) %>%
  dplyr::mutate(Genus_label_raw = shorten_genus(.data$Genus)) %>%
  dplyr::group_by(.data$Genus_label_raw) %>%
  dplyr::mutate(idx = dplyr::row_number(),
                Genus_label = dplyr::if_else(.data$idx == 1, .data$Genus_label_raw,
                                             paste0(.data$Genus_label_raw, ".", .data$idx))) %>%
  dplyr::ungroup() %>%
  dplyr::select(.data$Genus, .data$Genus_label)

## ============ 宽→长 + 合并分组 + 加显示标签 ============
long_df <- genus_rel_top %>%
  tidyr::pivot_longer(cols = dplyr::all_of(common_samples),
                      names_to = "Number", values_to = "RelAbund") %>%
  dplyr::left_join(meta, by = c("Number" = "Number")) %>%
  dplyr::left_join(label_map, by = "Genus") %>%
  dplyr::filter(!is.na(.data$Sampling_Location)) %>%
  dplyr::mutate(Sampling_Location = factor(.data$Sampling_Location, levels = loc_levels))

if (anyNA(long_df$Genus_label)) {
  idx <- is.na(long_df$Genus_label)
  long_df$Genus_label[idx] <- shorten_genus(long_df$Genus[idx])
}

## 列顺序（属标签）：按平均丰度
genus_order <- long_df %>%
  dplyr::group_by(.data$Genus_label) %>%
  dplyr::summarise(mean_abund = mean(.data$RelAbund, na.rm = TRUE), .groups = "drop") %>%
  dplyr::arrange(dplyr::desc(.data$mean_abund)) %>%
  dplyr::pull(.data$Genus_label)
long_df$Genus_label <- factor(long_df$Genus_label, levels = genus_order)

## 行顺序（样本）：分组内聚类
order_by_hclust <- function(df_one_group){
  w <- tidyr::pivot_wider(df_one_group,
                          id_cols = "Number",
                          names_from = "Genus_label",
                          values_from = "RelAbund",
                          values_fill = 0)
  mat <- as.matrix(w[, setdiff(colnames(w), "Number"), drop = FALSE])
  rownames(mat) <- w$Number
  if (nrow(mat) >= 2) {
    dd <- stats::dist(mat, method = "euclidean")
    hc <- stats::hclust(dd, method = "ward.D2")
    rownames(mat)[hc$order]
  } else rownames(mat)
}
sample_order_list <- lapply(levels(long_df$Sampling_Location), function(grp) {
  df_g <- long_df %>% dplyr::filter(.data$Sampling_Location == grp)
  if (nrow(df_g) == 0) return(character(0))
  order_by_hclust(df_g)
})
names(sample_order_list) <- levels(long_df$Sampling_Location)
chunks <- split(long_df, long_df$Sampling_Location, drop = TRUE)
for (grp in names(chunks)) {
  df <- chunks[[grp]]
  levs <- intersect(sample_order_list[[grp]], unique(df$Number))
  df$Number <- factor(df$Number, levels = levs)
  chunks[[grp]] <- df
}
long_df <- do.call(rbind, chunks)
long_df$Sampling_Location <- factor(long_df$Sampling_Location, levels = loc_levels)

## ============ 数值转换 + 分位数裁剪 ============
if (transform_mode == "log10") {
  raw_val <- log10(long_df$RelAbund + pseudo_count)
  fill_name <- paste0("log10(Rel.Abund.% + ", pseudo_count, ")")
} else {
  raw_val <- long_df$RelAbund
  fill_name <- "Relative abundance (%)"
}
stopifnot(length(raw_val) > 0)
if (is.null(heat_limits)) {
  qs <- stats::quantile(raw_val, probs = clip_quantiles, na.rm = TRUE)
  heat_limits <- c(qs[[1]], qs[[2]])
}
long_df$HeatVal <- pmin(pmax(raw_val, heat_limits[1]), heat_limits[2])

## ============ 顶部 Phylum 条带数据 & 自动补色 ============
col_phylum_df <- genus_rel_top %>%
  dplyr::distinct(.data$Genus, .data$Phylum) %>%
  dplyr::left_join(label_map, by = "Genus") %>%
  dplyr::select(.data$Genus_label, .data$Phylum) %>%
  dplyr::mutate(Genus_label = factor(.data$Genus_label, levels = levels(long_df$Genus_label)))

phyla_needed <- unique(col_phylum_df$Phylum)
phyla_needed <- phyla_needed[!is.na(phyla_needed) & phyla_needed != ""]

ensure_phylum_colors <- function(base_cols, needed, include_others_color = TRUE){
  missing <- setdiff(needed, names(base_cols))
  if (length(missing) > 0) {
    n <- length(missing)
    hues <- seq(15, 375, length.out = n + 1)[1:n]
    add_cols <- setNames(vapply(hues, function(h) grDevices::hcl(h = h, c = 70, l = 50), ""),
                         missing)
    base_cols <- c(base_cols, add_cols)
  }
  if (include_others_color && !"Others" %in% names(base_cols)) {
    base_cols <- c(base_cols, others_color)
  }
  base_cols
}
phylum_colors_all <- ensure_phylum_colors(phylum_colors_base, phyla_needed, include_others_color = TRUE)

phylum_breaks <- unique(col_phylum_df$Phylum)
if ("Others" %in% phylum_breaks) phylum_breaks <- c(setdiff(phylum_breaks, "Others"), "Others")
phylum_labels <- setNames(gsub("_.*", "", as.character(phylum_breaks)), phylum_breaks)
col_phylum_df$Phylum <- factor(col_phylum_df$Phylum, levels = phylum_breaks)

## ============ 组色渐变（白 → 组色） ============
mk_grad <- function(main_col, n = 7){ grDevices::colorRampPalette(c("#FFFFFF", main_col))(n) }
group_gradients <- lapply(group_main_colors, mk_grad)

## ============ 顶部：Phylum 条带（提 legend） ============
p_top_with_leg <- ggplot2::ggplot(col_phylum_df, ggplot2::aes(x = .data$Genus_label, y = 1, fill = .data$Phylum)) +
  ggplot2::geom_tile() +
  ggplot2::scale_fill_manual(values = phylum_colors_all[phylum_breaks],
                             breaks = phylum_breaks,
                             labels = phylum_labels,
                             drop   = TRUE,
                             name   = "Phylum") +
  ggplot2::coord_cartesian(clip = "off") +
  ggplot2::theme_void(base_size = 9) +
  ggplot2::theme(legend.position = "right", legend.title = ggplot2::element_text(face = "bold"),
                 plot.margin = ggplot2::margin(0, 5.5, 0, 5.5))
phylum_legend_grob <- cowplot::get_legend(p_top_with_leg)
p_top <- p_top_with_leg + ggplot2::theme(legend.position = "none")

## ============ 列聚类树（展示用，不改列顺序） ============
col_mat_df <- long_df %>%
  dplyr::select(dplyr::all_of(c("Number","Genus_label","HeatVal"))) %>%
  dplyr::distinct() %>%
  tidyr::pivot_wider(
    id_cols      = "Number",
    names_from   = "Genus_label",
    values_from  = "HeatVal",
    values_fill  = 0
  )
target_order <- levels(long_df$Genus_label)
col_mat <- as.matrix(col_mat_df[, target_order, drop = FALSE])

dist_cols <- stats::dist(t(col_mat), method = "euclidean")
hc_cols   <- stats::hclust(dist_cols, method = "ward.D2")
dend <- as.dendrogram(hc_cols) |> dendextend::rotate(order = target_order)
dd   <- ggdendro::dendro_data(dend, type = "rectangle")
seg  <- dd$segments
p_dend <- ggplot2::ggplot() +
  ggplot2::geom_segment(data = seg, ggplot2::aes(x = x, y = y, xend = xend, yend = yend), linewidth = 0.4) +
  ggplot2::scale_x_continuous(limits = c(0.5, length(target_order) + 0.5), expand = c(0, 0)) +
  ggplot2::scale_y_continuous(expand = c(0, 0)) +
  ggplot2::theme_void(base_size = 9) +
  ggplot2::theme(plot.margin = ggplot2::margin(0, 5.5, 0, 5.5))

## ============ 子图基础主题 & 绘制三组 ============
base_theme <- ggplot2::theme_bw(base_size = 9) +
  ggplot2::theme(panel.grid = ggplot2::element_blank(),
                 panel.border = ggplot2::element_rect(colour = "black", linewidth = 0.6),
                 axis.title = ggplot2::element_blank(),
                 axis.ticks.y = ggplot2::element_blank(),
                 axis.text.y  = ggplot2::element_blank(),
                 axis.ticks.x = ggplot2::element_blank(),
                 plot.margin  = ggplot2::margin(0, 5.5, 0, 10))

plot_one_group <- function(grp, show_x_text = FALSE, show_legend = FALSE,
                           y_title_angle = 90, x_angle = 90){
  df_g <- long_df %>% dplyr::filter(.data$Sampling_Location == grp)
  hjust_val <- if (y_title_angle == 0) 0 else 0.5
  ggplot2::ggplot(df_g, ggplot2::aes(x = .data$Genus_label, y = .data$Number, fill = .data$HeatVal)) +
    ggplot2::geom_tile() +
    ggplot2::scale_fill_gradientn(colours = group_gradients[[grp]], limits = heat_limits, name = grp) +
    base_theme +
    ggplot2::labs(y = grp) +
    ggplot2::theme(
      axis.title.y    = ggplot2::element_text(face = "bold", angle = y_title_angle,
                                              hjust = hjust_val, vjust = 0.5,
                                              margin = ggplot2::margin(r = 6)),
      axis.text.x     = if (show_x_text)
        ggplot2::element_text(angle = x_angle, hjust = 1, vjust = 1, size = 6)
      else ggplot2::element_blank(),
      legend.position = if (show_legend) "bottom" else "none",
      legend.title    = ggplot2::element_text(face = "bold"),
      legend.key.height = grid::unit(3.5, "mm")
    )
}

p_PH_main   <- plot_one_group("PH-NA", show_x_text = FALSE, show_legend = FALSE, y_title_angle = 90)
p_CPNA_main <- plot_one_group("CP-NA", show_x_text = FALSE, show_legend = FALSE, y_title_angle = 90)
p_CPAD_main <- plot_one_group("CP-AD", show_x_text =  TRUE, show_legend = FALSE,
                              y_title_angle = 90, x_angle = 45) +
  ggplot2::theme(plot.margin = ggplot2::margin(t = 0, r = 5.5, b = 18, l = 10))

## 取 3 条色条 → 横向一行 → 放在第 3 图下面
p_PH_legplot   <- plot_one_group("PH-NA", show_x_text = FALSE, show_legend = TRUE)  + ggplot2::theme(axis.text.x = ggplot2::element_blank(), legend.box.margin = ggplot2::margin(t = 6))
p_CPNA_legplot <- plot_one_group("CP-NA", show_x_text = FALSE, show_legend = TRUE) + ggplot2::theme(axis.text.x = ggplot2::element_blank(), legend.box.margin = ggplot2::margin(t = 6))
p_CPAD_legplot <- plot_one_group("CP-AD", show_x_text = FALSE, show_legend = TRUE) + ggplot2::theme(axis.text.x = ggplot2::element_blank(), legend.box.margin = ggplot2::margin(t = 6))
leg_PH   <- cowplot::get_legend(p_PH_legplot)
leg_CPNA <- cowplot::get_legend(p_CPNA_legplot)
leg_CPAD <- cowplot::get_legend(p_CPAD_legplot)
leg_row <- cowplot::plot_grid(cowplot::ggdraw(leg_PH), cowplot::ggdraw(leg_CPNA), cowplot::ggdraw(leg_CPAD),
                              nrow = 1, rel_widths = c(1,1,1))

p_CPAD_block <- p_CPAD_main / leg_row + patchwork::plot_layout(heights = c(10, 2.2))

## 体部分
p_body <- p_PH_main / p_CPNA_main / p_CPAD_block

## 左列堆叠：列树 / Phylum 条带 / 主体
p_left_stack <- p_dend / p_top / p_body + patchwork::plot_layout(heights = c(0.18, 0.06, 2.76))

## 右侧门图例与主体中间放空白列，拉开距离
p_spacer <- patchwork::plot_spacer()
p_leg    <- cowplot::ggdraw(phylum_legend_grob)

p_all <- p_left_stack | p_spacer | p_leg
p_all <- p_all + patchwork::plot_layout(widths = c(1, 0.10, 0.16))

## ============ 导出 ============
ggplot2::ggsave("perio_genus_heatmap_with_colDend_rightPhylumLegend_NC.pdf", p_all,
                width = fig_width_mm, height = fig_height_mm, units = "mm", bg = "transparent")
ggplot2::ggsave("perio_genus_heatmap_with_colDend_rightPhylumLegend_NC.svg", p_all,
                width = fig_width_mm, height = fig_height_mm, units = "mm", bg = "transparent")
ggplot2::ggsave("perio_genus_heatmap_with_colDend_rightPhylumLegend_NC.png", p_all,
                width = fig_width_mm, height = fig_height_mm, units = "mm", dpi = 600, bg = "transparent")

## （可选）调试：若仍出现 NA 属名，打印来源
if (FALSE && anyNA(long_df$Genus_label)) {
  message("❗ Genus_label 出现 NA，来源如下：")
  print(long_df %>% dplyr::filter(is.na(.data$Genus_label)) %>% dplyr::distinct(.data$Genus) %>% dplyr::arrange(.data$Genus))
}

## =========================
##  统计表格导出（Genus）
## =========================

## 0) 保障：必要对象存在
stopifnot(exists("genus_rel"), exists("genus_rel_top"),
          exists("common_samples"), exists("loc_levels"),
          exists("long_df"), exists("genus_phylum_map"))

## ---------- 1) 逐样本矩阵（ALL） ----------
# genus_rel: 列 = Genus + 样本列 + mean_abund
tbl_bySample_all <- genus_rel %>%
  dplyr::select(dplyr::all_of(c("Genus", common_samples))) %>%
  dplyr::left_join(genus_phylum_map, by = "Genus") %>%
  # 列顺序：Genus, Phylum, 样本...
  dplyr::relocate(.data$Phylum, .after = .data$Genus)

utils::write.csv(tbl_bySample_all,
                 file = "Table_genus_relative_abundance_bySample.csv",
                 row.names = FALSE)

## ---------- 2) 逐样本矩阵（Top-N，与热图一致） ----------
tbl_bySample_topN <- genus_rel_top %>%
  dplyr::select(dplyr::all_of(c("Genus", common_samples))) %>%
  dplyr::left_join(genus_phylum_map, by = "Genus") %>%
  dplyr::relocate(.data$Phylum, .after = .data$Genus)

utils::write.csv(tbl_bySample_topN,
                 file = "Table_genus_relative_abundance_bySample_topN.csv",
                 row.names = FALSE)

## ---------- 3) 各组汇总（mean, sd, median, IQR, prevalence） ----------
# 使用 long_df：列含 Genus, RelAbund, Sampling_Location（已对齐并筛到 Top-N）
# 如果你希望统计**全部属**，把 long_df 换成 all_long_df（见注释块）
long_top_df <- long_df %>%
  dplyr::select(.data$Genus, .data$RelAbund, .data$Sampling_Location)

# 计算 prevalence（>0 的样本占比）
tbl_group <- long_top_df %>%
  dplyr::group_by(.data$Genus, .data$Sampling_Location) %>%
  dplyr::summarise(
    n           = dplyr::n(),
    prevalence  = mean(.data$RelAbund > 0, na.rm = TRUE),         # 0~1
    mean        = mean(.data$RelAbund, na.rm = TRUE),
    sd          = stats::sd(.data$RelAbund, na.rm = TRUE),
    median      = stats::median(.data$RelAbund, na.rm = TRUE),
    q25         = as.numeric(stats::quantile(.data$RelAbund, 0.25, na.rm = TRUE, names = FALSE)),
    q75         = as.numeric(stats::quantile(.data$RelAbund, 0.75, na.rm = TRUE, names = FALSE)),
    IQR         = q75 - q25,
    .groups     = "drop"
  ) %>%
  dplyr::left_join(genus_phylum_map, by = "Genus") %>%
  dplyr::relocate(.data$Phylum, .after = .data$Genus) %>%
  dplyr::mutate(Sampling_Location = factor(.data$Sampling_Location, levels = loc_levels)) %>%
  dplyr::arrange(.data$Genus, .data$Sampling_Location)

utils::write.csv(tbl_group,
                 file = "Table_genus_relative_abundance_byGroup.csv",
                 row.names = FALSE)

## ---------- 4) 整体均值排序（ALL 样本） ----------
tbl_overall <- genus_rel %>%
  dplyr::transmute(
    Genus = .data$Genus,
    overall_mean = rowMeans(dplyr::across(dplyr::all_of(common_samples)), na.rm = TRUE)
  ) %>%
  dplyr::left_join(genus_phylum_map, by = "Genus") %>%
  dplyr::relocate(.data$Phylum, .before = .data$overall_mean) %>%
  dplyr::arrange(dplyr::desc(.data$overall_mean))

utils::write.csv(tbl_overall,
                 file = "Table_genus_relative_abundance_overall.csv",
                 row.names = FALSE)

## ---------- 5) 组间差异：Kruskal–Wallis + 两两 Wilcoxon（FDR） ----------
comparisons <- list(c("PH-NA","CP-NA"),
                    c("PH-NA","CP-AD"),
                    c("CP-NA","CP-AD"))

# a) Kruskal–Wallis across 3 groups（Top-N集合；如需 ALL，把 long_top_df 换成 all_long_df）
kw_df <- long_top_df %>%
  dplyr::group_by(.data$Genus) %>%
  dplyr::summarise(
    p_kw = tryCatch(
      stats::kruskal.test(.data$RelAbund ~ .data$Sampling_Location)$p.value,
      error = function(e) NA_real_
    ),
    .groups = "drop"
  ) %>%
  dplyr::mutate(p_kw_adj = stats::p.adjust(.data$p_kw, method = "BH")) %>%
  dplyr::left_join(genus_phylum_map, by = "Genus") %>%
  dplyr::relocate(.data$Phylum, .after = .data$Genus) %>%
  dplyr::arrange(.data$p_kw_adj)

# b) Pairwise Wilcoxon + FDR（每一对在“所有属”中统一做 BH）
pairwise_results <- lapply(comparisons, function(cc){
  gA <- cc[1]; gB <- cc[2]
  sub <- long_top_df %>% dplyr::filter(.data$Sampling_Location %in% c(gA, gB))
  # 先计算所有属的原始 p，再一次性 BH
  raw_p <- sub %>%
    dplyr::group_by(.data$Genus) %>%
    dplyr::summarise(
      p = tryCatch(
        stats::wilcox.test(.data$RelAbund[.data$Sampling_Location == gA],
                           .data$RelAbund[.data$Sampling_Location == gB],
                           exact = FALSE)$p.value,
        error = function(e) NA_real_
      ),
      med_A = stats::median(.data$RelAbund[.data$Sampling_Location == gA], na.rm = TRUE),
      med_B = stats::median(.data$RelAbund[.data$Sampling_Location == gB], na.rm = TRUE),
      .groups = "drop"
    ) %>%
    dplyr::mutate(
      p_adj = stats::p.adjust(.data$p, method = "BH"),
      log2FC = log2(.data$med_A + 1e-6) - log2(.data$med_B + 1e-6),  # >0: A高
      winner = dplyr::case_when(
        is.na(.data$med_A) | is.na(.data$med_B) ~ NA_character_,
        .data$med_A >= .data$med_B ~ gA,
        TRUE ~ gB
      ),
      Comparison = paste(gA, "vs", gB)
    ) %>%
    dplyr::left_join(genus_phylum_map, by = "Genus") %>%
    dplyr::relocate(.data$Phylum, .after = .data$Genus)
  raw_p
}) %>% dplyr::bind_rows() %>%
  dplyr::arrange(.data$Comparison, .data$p_adj, dplyr::desc(abs(.data$log2FC)))

# 合并 KW 与两两（方便一表引用）
kw_pair_merged <- pairwise_results %>%
  dplyr::left_join(kw_df %>% dplyr::select(.data$Genus, .data$p_kw, .data$p_kw_adj),
                   by = "Genus") %>%
  dplyr::relocate(.data$p_kw, .data$p_kw_adj, .after = .data$Phylum)

utils::write.csv(kw_pair_merged,
                 file = "Table_genus_KW_and_pairwise.csv",
                 row.names = FALSE)

## ---------- （可选）如果你想基于“全部属”而不是热图 Top-N 来做统计 ----------
## 取消以下注释，并将上面的 long_top_df 改为 all_long_df 即可
# all_long_df <- genus_rel %>%
#   dplyr::select(dplyr::all_of(c("Genus", common_samples))) %>%
#   tidyr::pivot_longer(cols = dplyr::all_of(common_samples),
#                       names_to = "Number", values_to = "RelAbund") %>%
#   dplyr::left_join(meta, by = c("Number" = "Number")) %>%
#   dplyr::filter(!is.na(.data$Sampling_Location)) %>%
#   dplyr::mutate(Sampling_Location = factor(.data$Sampling_Location, levels = loc_levels))
```
