## Supplementary material for "Niche-specific microbial community structure of subgingival plaque in periodontitis": periodontitis_reproducibility_PRJNA1512026: paired_design_reanalysis.py

```
"""Reanalyse alpha/beta diversity respecting the study's paired design.

PH-NA vs CP-NA is an independent comparison.  CP-NA vs CP-AD is a
within-participant comparison, with label swaps restricted within each pair.
"""
from pathlib import Path
import numpy as np
import pandas as pd
from scipy.spatial.distance import pdist, squareform
from scipy.stats import mannwhitneyu, wilcoxon

HERE = Path(__file__).resolve().parent
meta = pd.read_excel(HERE / "metadata.xlsx")
asv = pd.read_csv(HERE / "ASV_taxonomy_table.csv")
meta["Number"] = meta["Number"].astype(str)

def genus_abundance(sample_ids):
    g = asv["Genus"].fillna("Unclassified").astype(str).replace("", "Unclassified")
    counts = asv.loc[:, sample_ids].apply(pd.to_numeric, errors="coerce").fillna(0)
    counts.insert(0, "Genus", g)
    return counts.groupby("Genus", sort=False).sum().T.to_numpy(dtype=float)

def bray(x):
    return squareform(pdist(x, metric="braycurtis"))

def permanova_stat(d, labels):
    labels=np.asarray(labels); n=len(labels); groups=np.unique(labels)
    total=(d**2).sum()/(2*n)
    within=sum((d[np.ix_(labels==g,labels==g)]**2).sum()/(2*(labels==g).sum()) for g in groups)
    between=total-within
    return (between/(len(groups)-1))/(within/(n-len(groups))), between/total

def pcoa_coords(d):
    n=d.shape[0]; h=np.eye(n)-np.ones((n,n))/n
    b=-0.5*h@(d**2)@h
    vals, vecs=np.linalg.eigh(b); keep=vals>1e-10
    return vecs[:,keep]*np.sqrt(vals[keep])

def dispersion(d, labels):
    z=pcoa_coords(d); labels=np.asarray(labels)
    out=[]
    for g in np.unique(labels):
        idx=labels==g; centroid=z[idx].mean(axis=0)
        out.append(np.linalg.norm(z[idx]-centroid,axis=1))
    return out

def unpaired_result(name, ids_a, ids_b, nperm=999, seed=20260812):
    ids=list(ids_a)+list(ids_b); labels=np.array(["A"]*len(ids_a)+["B"]*len(ids_b)); d=bray(genus_abundance(ids))
    f,r2=permanova_stat(d,labels); rng=np.random.default_rng(seed)
    fp=np.array([permanova_stat(d,rng.permutation(labels))[0] for _ in range(nperm)])
    da,db=dispersion(d,labels)
    return {"comparison":name,"design":"independent","n_A":len(ids_a),"n_B":len(ids_b),"PERMANOVA_F":f,"PERMANOVA_R2":r2,"PERMANOVA_P":(1+(fp>=f).sum())/(nperm+1),"dispersion_test":"Mann–Whitney U","dispersion_P":mannwhitneyu(da,db,alternative="two-sided").pvalue}

def paired_result(name, pair_meta, nperm=999, seed=20260812):
    pair_meta=pair_meta.sort_values(["PatientID","Sampling_Location"])
    ids=pair_meta.Number.tolist(); labels=pair_meta.Sampling_Location.to_numpy(); d=bray(genus_abundance(ids))
    f,r2=permanova_stat(d,labels); rng=np.random.default_rng(seed); pids=pair_meta.PatientID.to_numpy()
    fp=[]
    for _ in range(nperm):
        lab=labels.copy()
        for pid in np.unique(pids):
            idx=np.where(pids==pid)[0]
            if rng.integers(2): lab[idx]=lab[idx][::-1]
        fp.append(permanova_stat(d,lab)[0])
    da,db=dispersion(d,labels)
    # In sorted rows, A/B distances must be aligned by participant.
    disp=pd.DataFrame({"PatientID":pids,"Sampling_Location":labels,"distance":np.concatenate([da,db])})
    # Construct from full coordinates to preserve row alignment.
    z=pcoa_coords(d); distances=[]
    for g in np.unique(labels):
        idx=labels==g; centroid=z[idx].mean(axis=0)
        distances.extend(zip(pids[idx], labels[idx], np.linalg.norm(z[idx]-centroid,axis=1)))
    disp=pd.DataFrame(distances,columns=["PatientID","Sampling_Location","distance"]).pivot(index="PatientID",columns="Sampling_Location",values="distance")
    return {"comparison":name,"design":"paired, labels permuted within participant","n_A":(labels=="CP-NA").sum(),"n_B":(labels=="CP-AD").sum(),"PERMANOVA_F":f,"PERMANOVA_R2":r2,"PERMANOVA_P":(1+(np.asarray(fp)>=f).sum())/(nperm+1),"dispersion_test":"Wilcoxon signed-rank","dispersion_P":wilcoxon(disp["CP-NA"],disp["CP-AD"]).pvalue}

healthy=meta.query("Sampling_Location == 'PH-NA'")
cpna=meta.query("Sampling_Location == 'CP-NA'")
cpad=meta.query("Sampling_Location == 'CP-AD'")
results=[unpaired_result("PH-NA vs CP-NA",healthy.Number,cpna.Number), paired_result("CP-NA vs CP-AD",pd.concat([cpna,cpad]))]
pd.DataFrame(results).to_csv(HERE / "paired_design_beta_diversity.tsv",sep="\t",index=False)

def alpha_tests(frame, value_column, index_name):
    """Run the same design-aware comparisons for a sample-level alpha index."""
    # Exported alpha tables include the same sample in two display facets.
    # Each sample must enter a design-aware test only once.
    values = frame.drop_duplicates("Number", keep="first").set_index("Number")[value_column]
    h = values.loc[healthy.Number].astype(float)
    na = values.loc[cpna.Number].astype(float)
    ad = values.loc[cpad.Number].astype(float)
    return pd.DataFrame([
        ["PH-NA vs CP-NA", "independent", len(h), len(na), h.mean(), na.mean(),
         mannwhitneyu(h, na, alternative="two-sided").pvalue],
        ["CP-NA vs CP-AD", "paired", len(na), len(ad), na.mean(), ad.mean(),
         wilcoxon(na.to_numpy(), ad.to_numpy()).pvalue],
    ], columns=["comparison", "design", "n_A", "n_B", f"mean_{index_name}_A",
                f"mean_{index_name}_B", "P_value"])

shannon = pd.read_csv(HERE / "Table_Alpha_Shannon_samples.csv")
shannon_out = alpha_tests(shannon, "Shannon", "Shannon")
shannon_out.to_csv(HERE / "paired_design_alpha_diversity.tsv", sep="\t", index=False)

chao1 = pd.read_csv(HERE / "Table_Chao1_samples.csv")
chao1_out = alpha_tests(chao1, "Value_Index", "Chao1")
chao1_out.to_csv(HERE / "paired_design_chao1_diversity.tsv", sep="\t", index=False)

print(shannon_out.to_string(index=False))
print(chao1_out.to_string(index=False))
print(pd.DataFrame(results).to_string(index=False))
```
