## Supplementary material for "Niche-specific microbial community structure of subgingival plaque in periodontitis": periodontitis_reproducibility_PRJNA1512026: phylogenetic_tree.R

```
## =========================================================
##  Genus 层面：每属取丰度最高序列 → 多序列比对 → 构树 → 作图
## =========================================================

suppressPackageStartupMessages({
  library(dplyr)
  library(tibble)
  library(stringr)
  library(Biostrings)
  library(DECIPHER)
  library(ape)
  library(phangorn)
  library(ggtree)
  library(ggplot2)
  library(scales)
})

## ---------------- 参数 ----------------
seqtab_rds <- "seqtab_nochim.rds"
tax_rds    <- "taxonomy.rds"

DROP_UNKNOWN_GENUS <- TRUE
ALIGN_ITER   <- 2
ALIGN_REFINE <- 2

## ---------------- 读入数据 ----------------
st   <- readRDS(seqtab_rds)
taxa <- readRDS(tax_rds)

st   <- as.matrix(st)
taxa <- as.data.frame(taxa, stringsAsFactors = FALSE)

seq_in_st   <- colnames(st)
seq_in_tax  <- rownames(taxa)
seq_common  <- intersect(seq_in_st, seq_in_tax)

if (length(seq_common) < 2) {
  stop("可匹配到的序列不足 2 条，无法构树。")
}

st_sub   <- st[, seq_common, drop = FALSE]
taxa_sub <- taxa[seq_common, , drop = FALSE]

## ---------------- 计算丰度并整理注释 ----------------
abund <- colSums(st_sub)
meta_seq <- tibble(
  Sequence = names(abund),
  TotalAbundance = as.numeric(abund),
  Genus  = if ("Genus"  %in% colnames(taxa_sub)) taxa_sub$Genus[match(names(abund), rownames(taxa_sub))] else NA_character_,
  Phylum = if ("Phylum" %in% colnames(taxa_sub)) taxa_sub$Phylum[match(names(abund), rownames(taxa_sub))] else NA_character_
)

# 清理 Genus/Phylum
meta_seq <- meta_seq %>%
  mutate(
    Genus  = ifelse(is.na(Genus)  | Genus  == "" , NA, Genus),
    Phylum = ifelse(is.na(Phylum) | Phylum == "" , "Unknown", Phylum)
  )

# 🔹 把 Phylum 名称中 "_" 及其后内容去掉
meta_seq <- meta_seq %>%
  mutate(Phylum = sub("_.*", "", Phylum))

if (DROP_UNKNOWN_GENUS) {
  n_before <- nrow(meta_seq)
  meta_seq <- meta_seq %>% filter(!is.na(Genus))
  n_after  <- nrow(meta_seq)
  message(sprintf("丢弃了 %d 条 Genus 缺失的序列；保留 %d 条用于属层面代表筛选。",
                  n_before - n_after, n_after))
} else {
  meta_seq <- meta_seq %>%
    mutate(Genus = ifelse(is.na(Genus), "Unknown_Genus", Genus))
}

## ---------------- 每个属选代表序列 ----------------
rep_by_genus <- meta_seq %>%
  group_by(Genus) %>%
  arrange(desc(TotalAbundance), .by_group = TRUE) %>%
  slice_head(n = 1) %>%
  ungroup()

dna <- DNAStringSet(rep_by_genus$Sequence)
tip_labels <- make.unique(rep_by_genus$Genus)
names(dna) <- tip_labels

tip_anno <- rep_by_genus %>%
  mutate(label = tip_labels) %>%
  select(label, Genus, Phylum, TotalAbundance)

## ---------------- 多序列比对 + 构树 ----------------
set.seed(1)
alignment <- AlignSeqs(dna, iterations = ALIGN_ITER, refinements = ALIGN_REFINE)
phydat <- phyDat(as(alignment, "matrix"), type = "DNA")
dm <- dist.ml(phydat)
treeNJ <- NJ(dm)

fit <- pml(treeNJ, data = phydat)
fitGTR <- update(fit, k = 4, inv = 0)
fitGTR <- optim.pml(
  fitGTR, model = "GTR",
  optInv = TRUE, optGamma = TRUE,
  optEdge = TRUE, optNni = TRUE,
  control = pml.control(trace = 0)
)
ml_tree <- fitGTR$tree
ml_tree$tip.label <- tip_anno$label

## ---------------- 作图 ----------------
tree_mid <- midpoint(ml_tree)
tree_mid <- ladderize(tree_mid, right = TRUE)
cap <- quantile(tree_mid$edge.length, 0.98, na.rm = TRUE)
tree_mid$edge.length <- pmin(tree_mid$edge.length, cap)

# 颜色映射：根据去掉后缀的门名
phyla <- sort(unique(tip_anno$Phylum))
pal <- hue_pal()(length(phyla))
names(pal) <- phyla
if ("Unknown" %in% names(pal)) pal["Unknown"] <- "grey70"

# Phylum 颜色方案（已去掉后缀）
phylum_colors <- c(
  "Absonditabacteria" = "#F39B7FFF",
  "Actinobacteria"    = "#E64B35FF",
  "Bacteroidetes"     = "#4DBBD5FF",
  "Cyanobacteria"     = "#00A087FF",
  "Firmicutes"        = "#3C5488FF",
  "Fusobacteria"      = "#8491B4FF",
  "Gracilibacteria"   = "#91D1C2FF",
  "Proteobacteria"    = "#0073C2FF",
  "Saccharibacteria"  = "#7E6148FF",
  "Spirochaetes"      = "#925E9FFF",
  "Synergistetes"     = "#FFD600FF",
  "Unknown"           = "grey70"
)

p <- ggtree(tree_mid, layout = "circular", size = 0.9, color = "black") %<+% tip_anno +
  geom_tippoint(aes(color = Phylum), size = 2, alpha = 0, show.legend = TRUE) +
  geom_tiplab2(
    aes(label = Genus, color = Phylum, angle = angle),
    size = 2.4, align = FALSE, linetype = 0, offset = 0.01,
    show.legend = FALSE
  ) +
  scale_color_manual(
    values = phylum_colors,
    guide = guide_legend(title = NULL, override.aes = list(shape = 16, size = 2, alpha = 1))
  ) +
  theme_tree() +
  theme(
    legend.position = "right",
    legend.margin = margin(l = 20),
    legend.box.margin = margin(5, 5, 5, 5),
    plot.title = element_text(hjust = 0.7, face = "bold", size = 14, margin = margin(b = 20)),
    plot.margin = margin(10, 40, 10, 10)
  ) +
  labs(title = "Genus phylogeny tree")

print(p)

## 导出
ggsave("phylo_genus_circular.png", p, width = 10, height = 10, dpi = 300)
ggsave("phylo_genus_circular.pdf", p, width = 10, height = 10)
ggsave("phylo_genus_circular.svg", p, width = 10, height = 10)
```
