## Supplementary material for "Niche-specific microbial community structure of subgingival plaque in periodontitis": periodontitis_reproducibility_PRJNA1512026: venn_diagram.R

```
## =========================================================
## 三组（牙周炎三水平：PH-NA、CP-NA、CP-AD）属级重叠分析 + Venn 图
## - 统计每组内各属的 ASV 数（richness）
## - 计算三组交集的属数
## - 绘制三集合 Venn（NC 风格配色）
## - 全程显式 dplyr/tidyr 命名空间，避免 select 冲突
## 依赖：readxl dplyr tidyr stringr ggplot2 ggvenn readr
## 输入：metadata.xlsx（含 Number, Sampling_Location）
##       ASV_taxonomy_table.csv（含 Genus + 各样本列，可有 ASV/ASV_ID）
## 输出：genus_asv_richness_by_group.csv
##       venn_genus_perio_3groups_NC.{pdf,svg,png}
##       （可选）genus_asv_richness_by_group_long.csv
## =========================================================

suppressPackageStartupMessages({
  library(readxl)
  library(dplyr)
  library(tidyr)
  library(stringr)
  library(ggplot2)
  library(ggvenn)
  library(readr)
})

## -------------------------
## 路径与输入文件
## -------------------------
meta_file   <- "metadata.xlsx"
asv_tax_csv <- "ASV_taxonomy_table.csv"

## 牙周炎三水平（固定顺序：健康在前）
loc_levels <- c("PH-NA","CP-NA","CP-AD")

## -------------------------
## 读取元数据（规范分组到三水平）
## -------------------------
clean_group <- function(x){
  y <- x %>%
    stringr::str_trim() %>% stringr::str_replace_all("\\s+","") %>%
    toupper() %>% stringr::str_replace_all("_","-")
  dplyr::case_when(
    y %in% c("PH-NA","PHNA","H-NA","HNA","HEALTHY","HEALTHYNA","PH") ~ "PH-NA",
    y %in% c("CP-NA","CPNA","NONATTACHED","NASP","N-SUB","NSUB")      ~ "CP-NA",
    y %in% c("CP-AD","CPAD","ATTACHED","ASP","A-SUB","ASUB")          ~ "CP-AD",
    TRUE ~ NA_character_
  )
}

meta <- readxl::read_xlsx(meta_file) %>%
  dplyr::mutate(Number = as.character(.data$Number),
                Sampling_Location = clean_group(.data$Sampling_Location)) %>%
  dplyr::select(dplyr::all_of(c("Number","Sampling_Location"))) %>%
  dplyr::filter(!is.na(.data$Number), !is.na(.data$Sampling_Location)) %>%
  dplyr::filter(.data$Sampling_Location %in% loc_levels) %>%
  dplyr::mutate(Sampling_Location = factor(.data$Sampling_Location, levels = loc_levels))

if (nrow(meta) == 0) stop("metadata 中无 PH-NA/CP-NA/CP-AD 记录。")

## -------------------------
## 读取 ASV × taxonomy 表
## -------------------------
asv_tab <- utils::read.csv(asv_tax_csv, check.names = FALSE, stringsAsFactors = FALSE)

## 样本列（与 meta$Number 交集）
sample_cols <- intersect(colnames(asv_tab), meta$Number)
if (length(sample_cols) == 0) {
  stop("在 ASV_taxonomy_table.csv 中没有找到与 metadata$Number 匹配的样本列。")
}

## Genus 列健壮性
if (!"Genus" %in% colnames(asv_tab)) {
  stop("ASV_taxonomy_table.csv 中未找到列 'Genus'。")
}
asv_tab$Genus <- as.character(asv_tab$Genus)
asv_tab$Genus[is.na(asv_tab$Genus) | asv_tab$Genus == ""] <- "Unassigned"

## ASV ID（若已有 ASV/ASV_ID 列则沿用）
if ("ASV_ID" %in% names(asv_tab)) {
  asv_tab$ASV_ID <- as.character(asv_tab$ASV_ID)
} else if ("ASV" %in% names(asv_tab)) {
  asv_tab$ASV_ID <- as.character(asv_tab$ASV)
} else {
  asv_tab$ASV_ID <- paste0("ASV_", seq_len(nrow(asv_tab)))
}

## -------------------------
## 转长表 + 合并分组（>0 视为出现）
## -------------------------
long_df <- asv_tab %>%
  dplyr::select(dplyr::all_of(c("ASV_ID","Genus", sample_cols))) %>%
  tidyr::pivot_longer(cols = dplyr::all_of(sample_cols),
                      names_to = "Number", values_to = "Count") %>%
  dplyr::mutate(Number = as.character(.data$Number)) %>%
  dplyr::left_join(meta, by = "Number") %>%
  dplyr::filter(!is.na(.data$Sampling_Location)) %>%
  dplyr::mutate(
    Count = suppressWarnings(as.numeric(.data$Count)),
    Count = dplyr::if_else(is.na(.data$Count), 0, .data$Count),
    Present = .data$Count > 0
  )

## -------------------------
## 1) 每个属在每组的 ASV 数（richness）
## -------------------------
groups <- loc_levels

genus_asv_richness <- long_df %>%
  dplyr::filter(.data$Present) %>%
  dplyr::group_by(.data$Genus, .data$Sampling_Location) %>%
  dplyr::summarise(ASV_n = dplyr::n_distinct(.data$ASV_ID), .groups = "drop") %>%
  tidyr::pivot_wider(
    names_from  = "Sampling_Location",
    values_from = "ASV_n",
    values_fill = 0
  ) %>%
  { df <- .;
  missing_cols <- setdiff(groups, names(df))
  if (length(missing_cols) > 0) df[missing_cols] <- 0
  df
  } %>%
  dplyr::mutate(
    Groups_present = rowSums(dplyr::across(dplyr::all_of(groups), ~ (.x > 0)), na.rm = TRUE),
    Shared_all3    = .data$Groups_present == 3
  ) %>%
  dplyr::arrange(dplyr::desc(.data$Groups_present), .data$Genus)

readr::write_csv(genus_asv_richness, "genus_asv_richness_by_group.csv")

## -------------------------
## 2) 三组共同属数量（交集大小）
## -------------------------
genus_sets <- long_df %>%
  dplyr::filter(.data$Present) %>%
  dplyr::group_by(.data$Sampling_Location, .data$Genus) %>%
  dplyr::summarise(has = any(.data$Present), .groups = "drop") %>%
  dplyr::filter(.data$has) %>%
  dplyr::group_by(.data$Sampling_Location) %>%
  dplyr::summarise(Genus_set = list(sort(unique(.data$Genus))), .groups = "drop")

get_set <- function(g) {
  x <- genus_sets$Genus_set[genus_sets$Sampling_Location == g]
  if (length(x) == 0) character(0) else x[[1]]
}
set_PHNA <- get_set("PH-NA")
set_CPNA <- get_set("CP-NA")
set_CPAD <- get_set("CP-AD")

shared_all3 <- length(Reduce(intersect, list(set_PHNA, set_CPNA, set_CPAD)))
cat("三组共同具有的属数量（交集）：", shared_all3, "\n")

## -------------------------
## 3) 三集合 Venn 图（NC 风）
##    颜色顺序需与 set_list 顺序一致
## -------------------------
set_list <- list(
  "PH-NA" = set_PHNA,
  "CP-NA" = set_CPNA,
  "CP-AD" = set_CPAD
)

p_venn <- ggvenn::ggvenn(
  set_list,
  fill_color    = c("#009E73", "#0072B2", "#CC79A7"),  # PH-NA, CP-NA, CP-AD
  stroke_size   = 0.8,
  set_name_size = 4.2,
  text_size     = 4.2,
  show_percentage = FALSE
) +
  ggplot2::theme_void() +
  ggplot2::theme(
    legend.position = "none",
    plot.margin = ggplot2::margin(5.5, 5.5, 5.5, 5.5),
    plot.background = ggplot2::element_rect(fill = NA, colour = NA)
  ) +
  ggplot2::labs(title = "Genera overlap among periodontal groups ") +
  ggplot2::theme(
    plot.title = ggplot2::element_text(hjust = 0.5, size = 14, face = "bold"),
    plot.title.position = "panel",
    plot.margin = ggplot2::margin(8, 6, 6, 6)
  )

ggplot2::ggsave("venn_genus_perio_3groups_NC.pdf", p_venn,
                width = 160, height =120, units = "mm",
                device = cairo_pdf, bg = "transparent")
ggplot2::ggsave("venn_genus_perio_3groups_NC.png", p_venn,
                width = 160, height = 120, units = "mm",
                dpi = 600, bg = "transparent")
ggplot2::ggsave("venn_genus_perio_3groups_NC.svg", p_venn,
                width = 160, height = 120, units = "mm",
                bg = "transparent")

## -------------------------
## （可选）导出“每组每属 ASV 数”的长表
## -------------------------
genus_asv_richness_long <- long_df %>%
  dplyr::filter(.data$Present) %>%
  dplyr::group_by(.data$Genus, .data$Sampling_Location) %>%
  dplyr::summarise(ASV_n = dplyr::n_distinct(.data$ASV_ID), .groups = "drop") %>%
  dplyr::arrange(.data$Genus, .data$Sampling_Location)

# 如需另存：取消下一行注释
# readr::write_csv(genus_asv_richness_long, "genus_asv_richness_by_group_long.csv")
```
