## Supplementary material for "Niche-specific microbial community structure of subgingival plaque in periodontitis": periodontitis_reproducibility_PRJNA1512026: README.md

# Reproducibility archive — PRJNA1512026

This archive supports the preprint *Niche-specific microbial community structure of subgingival plaque in periodontitis*.

## Study design

The de-identified analysis set contains 82 samples: 36 paired non-attached and attached subgingival plaque sample pairs from participants with periodontitis (CP-NA and CP-AD) and 10 non-attached plaque samples from healthy controls (PH-NA).

## Data

Raw 16S rRNA V3–V4 reads are being submitted to NCBI SRA under BioProject PRJNA1512026. The BioProject and 82 BioSample records have been processed. Individual SRA run and BioSample accessions will be added after SRA processing and public release.

`inputs/` contains de-identified processed analysis inputs, not raw sequencing reads. `outputs/` contains the design-aware diversity statistics reported in Supplementary Table S1. `code/` contains the analysis scripts supplied as archived research code.

The association-network code uses Spearman correlations after within-group prevalence filtering; retained edges meet |rho| >= 0.5 and Benjamini–Hochberg-adjusted P < 0.05. The retained edge tables are included in `outputs/`.

## Software

Primary processing: DADA2 v1.28.0 in R v4.3.2; SILVA release 138.1. Python analysis requires NumPy, pandas, SciPy, and openpyxl. R scripts declare their required packages.

## Limits

The code is provided to document and support reproduction of the reported analyses. Raw reads and individual accession mappings are obtained through NCBI SRA after public release. No direct participant identifiers are included.
